# Experimental evolution of root-associated microbial communities

**DOI:** 10.64898/2026.09.18.752712

**Authors:** Niklas Kiel, Kathrin Wippel, Ruben Garrido-Oter

## Abstract

Microbial communities associated with plants are dynamic ecosystems whose structure and functions influence host health and fitness. While the ecological roles and beneficial function of plant microbiomes are increasingly well understood, the evolutionary dynamics that shape them over time remain unclear. Experimental evolution constitutes a powerful tool to study microbial adaptation, but so far, its application has been largely focused on free-living or single-species populations. Here, we use experimental evolution to test whether taxonomically complex synthetic bacterial communities can adapt to novel plant hosts. We show that replicate microbial communities undergo rapid and parallel genomic evolution, with measurable gains in colonization fitness. Adaptive changes involved mutations in regulatory processes, motility, and core metabolism, leading to significant fitness gains across diverse taxa. Our findings demonstrate that the plant host can drive microbiome evolution without disrupting the stability of the root microbiota.

## Introduction

Plants live in close association with communities of diverse microbes that can impact their health, growth and fitness^1–4^. These communities can provide beneficial functions such as nutrient mobilization^5^, protection against pathogens^6–9^, and promote tolerance to environmental stresses^10^. Root-associated microbiota, which originate from a small fraction of the vastly diverse soil biome and assemble within days^11,12^, have emerged as crucial determinants of plant performance. Collectively, these microbiota are increasingly viewed as integral components of plant adaptive strategies to environmental factors, such as nutrient limitation^13^ or drought^14^.

Host factors impose selective pressures on members of these communities, potentially driving evolutionary processes that shape community composition, function, and host specificity over ecological and evolutionary timescales^15^. While the ecology and functions of plant microbiota are increasingly well understood, their capacity to evolve and adapt to different host species remains largely unexplored. Recent work has shown that individual members of the plant microbiome can adapt rapidly to novel host environments, leading to changes in colonization efficiency and functional outcomes^16–18^. However, how such evolutionary change unfolds in a community context, where selection acts in the presence of interspecific interactions and shifting community structure, remains poorly understood.

Experimental evolution offers a powerful framework to study microbial adaptation, enabling controlled tracking of evolutionary trajectories at high resolution over short timescales^19,20^. Long-term evolution experiments (LTEEs), in particular, have shown that evolutionary responses can be highly reproducible across replicate populations exposed to the same selective pressures^21–23^. In bacteria, LTEEs have revealed convergent patterns of mutation and sustained fitness gains over tens of thousands of generations^22,24,25^. Similar findings have been observed in LTEEs with eukaryotes such as *Saccharomyces cerevisiae*, where evolutionary outcomes appear to be the result of both chance and selection^26^. Yet, in natural systems, microbes evolve within communities, where interspecific interactions can shape both ecological dynamics and adaptive trajectories. Several studies have begun to address this complexity by experimentally evolving simplified communities comprising two or more species, revealing that such interactions can constrain or redirect evolution, and alter functional outputs^27–34^. Yet most of these studies have focused on free-living communities with limited taxonomic complexity, and few have examined evolutionary dynamics in host-associated microbiomes.

In previous work^35^, we showed that the root microbiota of the brassica *Arabidopsis thaliana* (*At*) and the legume *Lotus japonicus* (*Lj*), which diverged 125 million years ago^36^, display signatures of host adaptation, preferentially colonizing their native over non-native host species. This host preference phenotype suggested that members of the root microbiota undergo host species-specific adaptation. However, the genetic mechanisms underlying this process remain unresolved. Here, we sought to investigate how such adaptation occurs and whether host-imposed selective pressures can reshape microbial communities on timescales accessible to experimental manipulation. We hypothesized that repeated exposure to a novel plant species would impose strong selective pressures on these communities, driving adaptive genomic changes to improve colonization fitness on the new host.

In multi-species microbiota, fitness of individual species is not only determined by their capacity to colonize their host but also by competition with other community members for host-derived resources. We hypothesized that evolution within a taxonomically diverse community would reveal constraints and trade-offs absent in single-strain systems, for example, between optimizing host colonization and maintaining competitive capacity. By tracking how multiple interacting taxa respond to host-imposed selection, we aimed to identify generalizable patterns of adaptation that emerge in the structured environment of the root microbiome.

To test these hypotheses, we designed an *in planta* microbial evolution experiment using synthetic bacterial communities (SynComs), each passaged through both native and non-native host species over multiple plant generations. This framework allowed us to monitor evolutionary change across multiple bacterial taxa within a taxonomically diverse yet simplified host-associated microbiome. By tracking genomic and phenotypic changes over time, we sought to uncover the genetic basis of host adaptation and assess whether microbiome evolution follows predictable trajectories under host-specific selective pressures.

## Results

### Changes in community composition during experimental evolution

To determine whether plant hosts drive evolutionary change in their associated microbiota, we conducted an evolution experiment using two bacterial SynComs: *At*-SC, derived from *Arabidopsis thaliana*^37^, and *Lj*-SC, derived from *Lotus japonicus*^35^. For balanced diversity, these two SynComs were designed to include one representative bacterial strain from each taxonomic family present in the *At* and *Lj* microbial culture collections^35,37^. We repeatedly passaged them through roots of either their native or non-native plant host using a soil-based gnotobiotic system^38^ (**Methods**; **Figure 1A**). This yielded four experimental conditions representing all combinations of SynCom and host species (**Figure 1B**): *At*-SC on *Arabidopsis* (*At*-SC*^At^*), *At*-SC on *Lotus* (*At*-SC*^Lj^*), *Lj*-SC on *Arabidopsis* (*Lj*-SC*^At^*), and *Lj*-SC on *Lotus* (*Lj*-SC*^Lj^*). Each cycle comprised 5 weeks of plant growth, after which root bacterial communities were extracted and used to inoculate the axenic roots of the subsequent plant generation. This process was repeated for 16 plant cycles (**Figure 1A**), which corresponds to an estimated ∼2,500 bacterial generations, on average across taxa (**Table S1**). At each harvesting step, we measured plant growth, extracted DNA samples for sequencing, and performed limiting dilutions to quantify absolute microbial abundances and isolate evolved bacterial strains (**Methods**).

**Figure 1.**
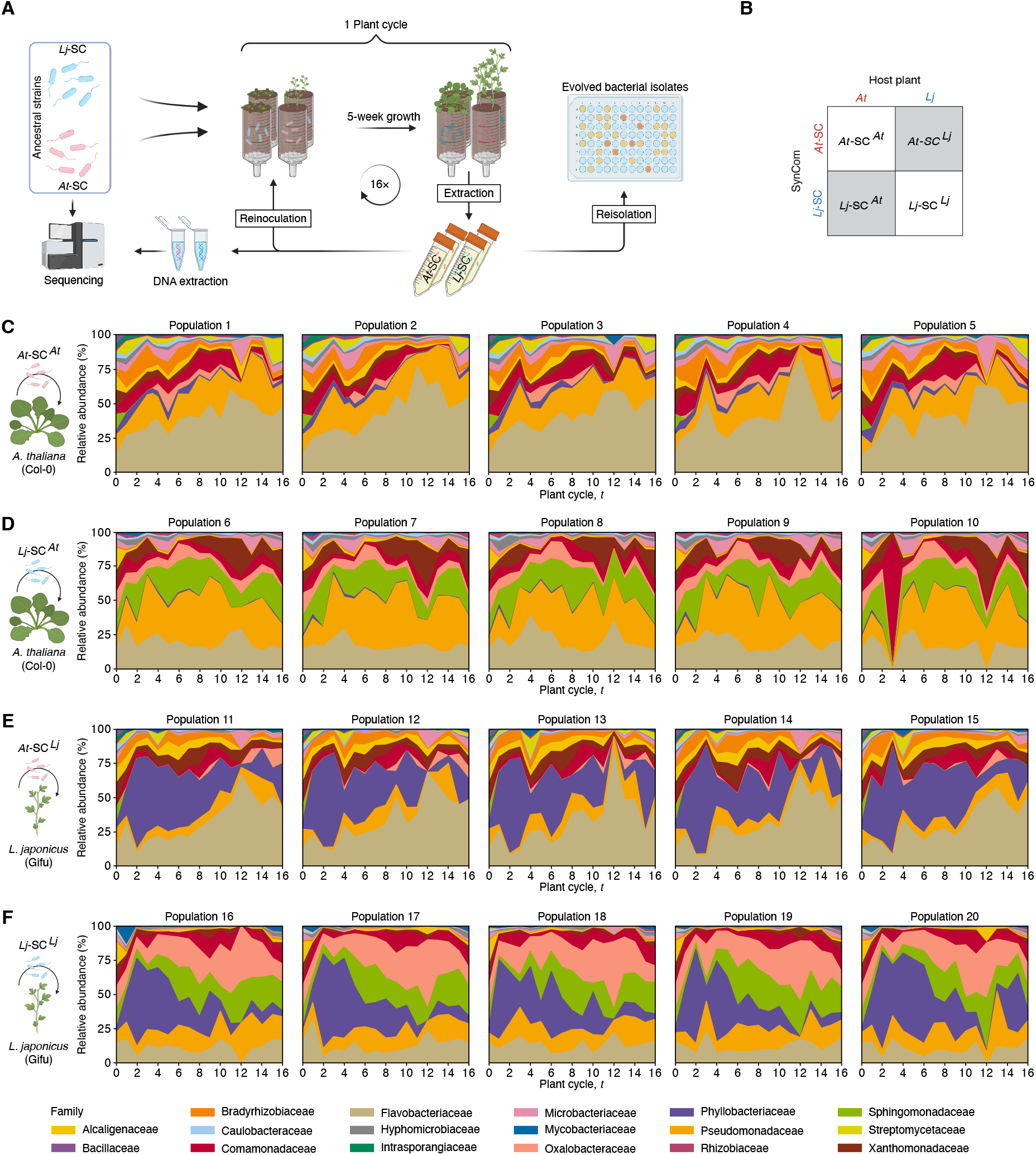
Experimental evolution of root-associated microbial communities. (**A**) Schematic overview of the conducted evolution experiment. (**B**) Summary of experimental conditions (2 plant species × 2 SynComs). Non-native conditions are indicated by dark grey. (**C**-**F**) Variation in community structure over time for SynComs *At*-SC*^At^* (**C**), *Lj*-SC*^At^*(**D**), *At*-SC*^Lj^* (**E**), and *Lj*-SC*^Lj^*(**F**). Bacterial abundances are shown as relative abundances (*y*-axis) plotted against time (*x*-axis). Different colours indicate the taxonomy of the respective strain.

Throughout the experiment, plant growth remained relatively constant, indicating stability in host-microbiota associations over time (**Figure S1A-S1B**). Neither pathogenic nor mutualistic interactions emerged from these repeated host passages. SynCom-inoculated plants showed no growth detriment compared to axenic conditions, and functional root nodule symbiosis remained intact in *Lotus* across all plant generations. Absolute bacterial abundances, assessed through colony-forming unit (CFU) counts from reisolated cultures (**Methods**), showed only a marginal increase throughout the experiment (**Figure S1C**). These observations may suggest that adaptation at the community level occurred without substantial expansion of ecological niches or increases in bacterial carrying capacity on either host.

Community profiling *via* 16S rRNA gene amplicon sequencing showed that bacterial diversity remained high across all treatments and time points, displaying complex patterns of community composition (**Figure 1C-1F**). Alpha diversity displayed no significant decline throughout the experiment (**Figure S2**). Most of the initial strains present in the start inocula (29 out of 34) persisted through all plant generations, highlighting remarkable community resilience. The exceptions included the two members of the Bacillaceae (Root131 and LjRoot5) and Intrasporangiaceae (Root101 and LjRoot24) families, and the *Lotus*-derived Streptomycetaceae (LjRoot303) strain, which were lost at different plant cycles (**Table S1**).

Comparison of community profiles across timepoints showed striking reproducibility among independent replicate populations, suggesting deterministic community assembly processes. The primary source of compositional variation was the host species, which accounted for a significant fraction of the total variance in microbial community diversity (PERMANOVA on Bray-Curtis dissimilarities; 28.43% of variance; *P* = 0.0001; **Figure 1C-1F**). Despite overall stability, relative abundances of specific taxa fluctuated over time, leading to shifts in community composition (19.06% of variance; *P* = 0.0001; **Figure 1C-1F**). Notably, one member of the Flavobacteriaceae family (Root935) increased in relative abundance (RA) by 90.1%, on average, from generation 1 to generation 16. Such dynamics could reflect evolutionary adaptation of individual strains to host-associated environments, resulting in fitness advantages over other community members. Alternatively, these changes over time may be the consequence of ecological interactions within the community, including competition and priority effects.

### Accumulation and fixation of genomic variants in root-associated bacterial communities

We hypothesized that throughout the 20-month evolution experiment, individual strains within the SynComs would accumulate *de novo* mutations in their genomes. According to the principle of natural selection, we expected that mutations conferring a fitness benefit would be positively selected and eventually reach fixation. Furthermore, we speculated that these genetic adaptations would impact competitive dynamics, potentially driving the measurable shifts in community composition reported in the preceding section. To investigate this, we performed shotgun sequencing of bacterial populations at multiple timepoints throughout our evolution experiment. Following stringent quality control, variant calling, and allele frequency estimation (**Methods**), we identified single-nucleotide variants (SNVs) and short insertions and deletions (indels) that arose and persisted in the genomes of all SynCom members over the course of the experiment. We observed consistent accumulation of these genetic variants in bacterial populations of both SynComs (*At*-SC and *Lj*-SC) in all experimental conditions, with a sustained increase over successive plant generations (**Figure 2A-B**). Both host environments (*Arabidopsis* and *Lotus* roots) drove comparable patterns of variant accumulation, which emerged consistently across independent replicate populations (**Figure 2C-2D**).

**Figure 2.**
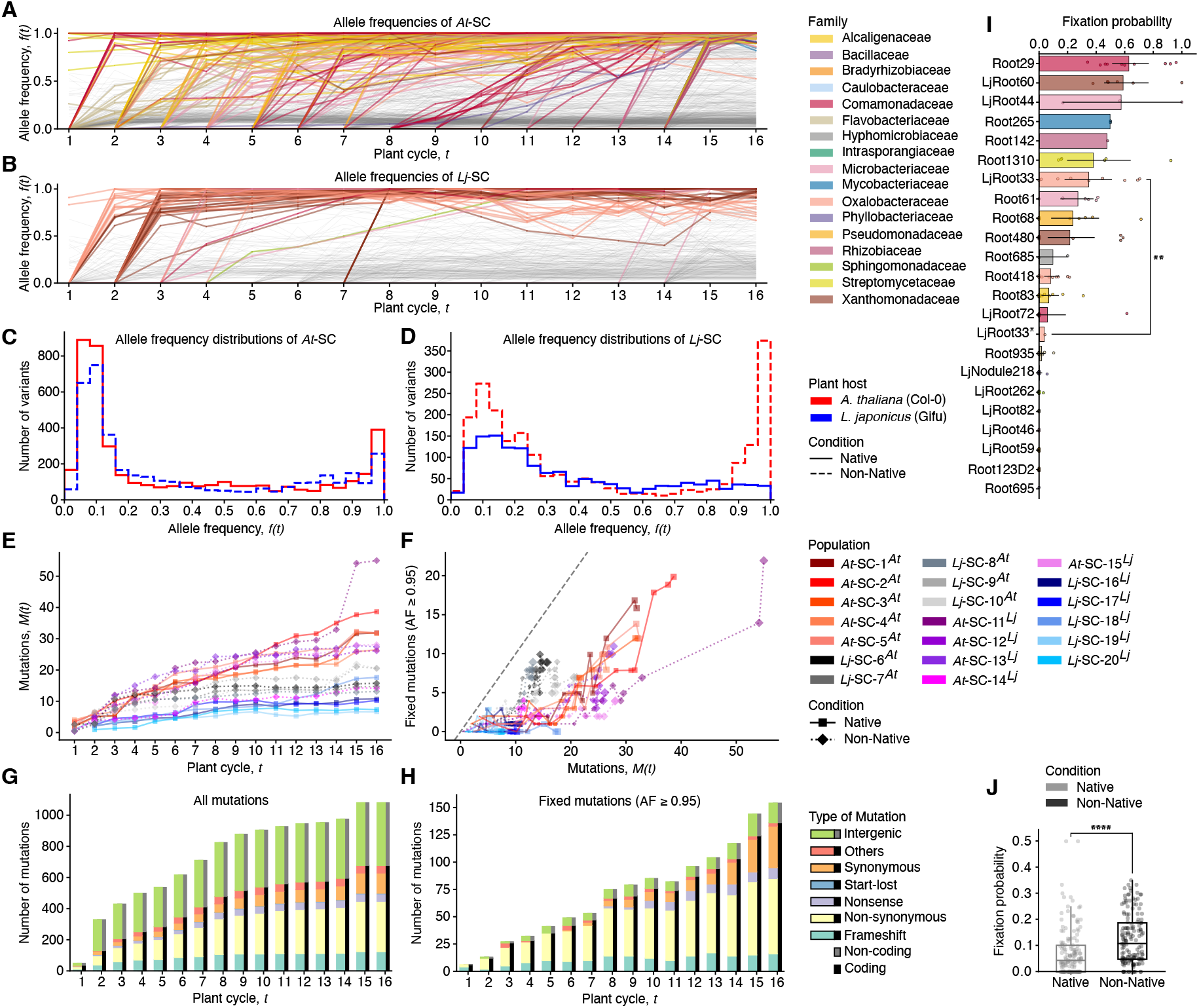
Population-level genetic diversity of evolving SynComs. Individual variant trajectories over time that occurred in evolving *At*-SC (**A**; *n* = 736) and *Lj*-SC (**B**; *n* = 344). Each line represents the allele frequency as a function of time of mutations that arose during the evolution experiment. Mutations that were fixed in the population (AF ≥ 0.95) are coloured according to the taxonomy of the strain. Non-fixed variants are depicted in grey. (**C**-**D**) Histograms of allele frequency distributions for *At*-SC (**C**), and *Lj*-SC (**D**). (**E**) Total derived allele frequency *M*(*t*) versus time coloured by replicate population. (**F**) Number of fixed mutations plotted against the *M*(*t*). (**G**-**H**) Number of mutations by class plotted against the plant cycle (*t*) for all and fixed variants (allele frequency ≥ 0.95), respectively. Different colours indicate different classes of mutations. Features affecting coding sequences are additionally coloured in black. Intergenic mutations are coloured in grey. (**I**) Strain-specific fixation probabilities calculated from the fraction of fixed mutations and all mutations (*n* = 151). Colours indicate taxonomy and are the same as in **A**-**B**. Comparison of LjRoot33 and the hypermutator was performed using a one-sample *t*-test. (**J**) Fixation probabilities between native and non-natively evolving SynComs (*n* = 313). Groups were compared using a Mann-Whitney *U* test.

Next, we calculated the rate at which mutations accumulate through the experiment, by calculating the total derived allele frequency *M*(*t*), defined as the sum of allele frequencies for all mutations observed in a population at a given time *t* (**Methods**). These values, which correspond to the rate of molecular evolution in a population, remained high but stable throughout the experiment (**Figure 2E**). A notable exception occurred in one replicate population, where the *Lotus*-derived Oxalobacteraceae strain (LjRoot33) acquired a mutation in the *mutS* mismatch repair gene, conferring a hypermutator phenotype^39^. This mutation drastically increased the rate of variant accumulation in this lineage by 158.57%, highlighting the emergence of hypermutability as a potential evolutionary strategy within host-associated communities.

However, despite overall similarity across conditions, considerable differences emerged when comparing variant accumulation rates among individual community members (**Figure S3**). For instance, strains from the Flavobacteriaceae family showed particularly high substitution rates (4.12 × 10^-6^ mutations × nucleotide^-1^ per generation, on average), whereas strains from other families such as Bradyrhizobiaceae accumulated significantly fewer mutations (4.19×10^-7^ mutations × nucleotide^-1^ × generation^-1^, on average). Interestingly, we observed a significant correlation between changes in relative abundance during the evolution experiment (**Figure 1C-1F**) and the total derived allele frequency *M*(*t*) per strain (**Figure S4A**). This suggests that the observed changes in community composition could be at least in part explained by adaptive evolution.

Analysis of allele frequencies over time indicated that a subset of these variants consistently reached fixation in a population, suggesting their significance for adaptation to the plant root environment (**Figure 2A-2B**). Conversely, another subset of variants persisted in replicate populations at lower abundances during multiple plant cycles, resulting in bi-modal distributions of allele frequencies (**Figure 2C-2D**). These patterns are consistent with clonal interference, where beneficial mutations compete within the population and only the most advantageous variants reach fixation. In this scenario, some beneficial mutations may not be fixed in the population if they are outcompeted by other bacterial lineages carrying alternative mutations with stronger fitness benefits. However, theory predicts that selective sweeps driven by the fittest lineage will eliminate competing beneficial mutations present in other lineages. Because each sweep fixes one allele to near 100% frequency, each fixed variant contributes approximately 1 to *M*(*t*), such that *M*(*t*) approximates the cumulative count of fixed variants^19^. Analysis of replicate lines from our evolution experiment displayed patterns consistent with this expectation (**Figure 2F**), indicating the importance of clonal interference as a driver of molecular evolution in bacterial populations, also in a community context.

We then tested if the observed genomic changes led to altered growth rates of individual community members. We employed a metagenomics-based method for estimating growth rates *in situ*, based on differences in read coverage distribution between the origin and terminus of replication in each bacterial genome^40,41^. We validated this approach by comparing *in situ* growth estimates with independently measured *in vitro* growth rates of ancestral strains, showing that these two metrics were significantly correlated (**Figure S4B**). Throughout the experiment, growth rates at the strain level remained stable, suggesting that the observed genomic changes did not lead to increased growth at the sampled timepoints (**Figure S4C**). Given that samples were collected after five weeks of *in planta* bacterial growth in each cycle, it is possible that significant changes in growth rates could only be observed during the initial stages of community establishment. Alternatively, adaptive mutations may primarily enhance other traits such as competitiveness within the community, or host colonization efficiency.

Further examination revealed only a weak correlation between strain-specific growth rates and the number of detected variants (**Figure S4D**). For instance, the two *Pseudomonas* strains (Root68 and LjRoot59) showed a high growth rate *in planta* but comparatively low substitution rate, while the Flavobacteriaceae strains (Root935 and LjRoot82) displayed the opposite pattern. These contrasting observations suggest that variation in substitution rates is not solely explained by growth dynamics, and that distinct selective pressures acting on different community members may also contribute to the observed patterns of variant accumulation.

### Fixed variants are likely adaptive and vary in occurrence between community members

To better understand the mechanisms underpinning bacterial adaptation in our evolution experiment, we examined what types of mutations appear and are fixed in replicate bacterial populations. The genomic distribution of variants was skewed toward coding regions, with 58.45% of all detected mutations located within protein-coding genes (**Figure 2G**). This pattern was more pronounced among fixed variants (allele frequency ≥ 95%), of which 88.68% were intragenic (**Figure 2H**). Further classification of these mutations also revealed that fixed variants exhibited a significantly elevated ratio of non-synonymous to synonymous substitutions compared to variants with lower allele frequencies (2.18-fold increase; *P* = 0.04). These data are consistent with positive selection acting preferentially on non-synonymous, likely functional, mutations.

We also observed considerable variation in fixation probability across individual bacterial strains (**Figure 2I**), ranging from approximately 60-80% for members of the Comamonadaceae family (LjRoot72 and Root29) to less than 10% in other lineages, such as Flavobacteriaceae (Root935 and LjRoot82). Interestingly, we observed a significant difference between the fixation probabilities of the hypermutator population of strain LjRoot33 (*Oxalobacteraceae*) and their non-mutator counterparts (4.05% *versus* 35.06%, *P* = 0.01). This pattern was expected given the higher rate at which hitchhiker mutations accumulate in hypermutator populations and is consistent with observations from previous LTEEs^22,25,42–45^.

In general, the observed differences in fixation probabilities between bacterial strains are unlikely the consequence of varying sequencing coverage, which we have accounted for in our analyses (**Methods**). Instead, it likely reflects differences in effective population size, mutation rate, or in the strength of the selective pressure imposed on each community member. Intriguingly, fixation probabilities were significantly higher in populations evolving on a non-native host species compared to those maintained on their native host (**Figure 2J**). This result supports the hypothesis of an increased evolutionary potential, or accelerated evolutionary change, when bacterial communities encounter new ecological contexts, such as a new host species.

### Parallel evolution of SynComs and reproducible genomic signatures of host adaptation

To further characterize genomic changes associated with bacterial adaptation to plant roots, we analysed the variant profiles using principal component analysis (PCA) based on allele frequencies of detected variants across experimental conditions and timepoints. Notably, replicate populations evolving independently under the same conditions (i.e., host species) exhibited remarkable similarity in their genomic trajectories, evidenced by their clustering in the PCA plots (**Figure 3A-3B**). Within each plant host species, samples also grouped by their plant generation, further illustrating the cumulative nature of genetic divergence and variant accumulation over time. These patterns suggest that repeated exposure to the same selective environment drives reproducible evolutionary trajectories, even in complex multispecies communities.

**Figure 3.**
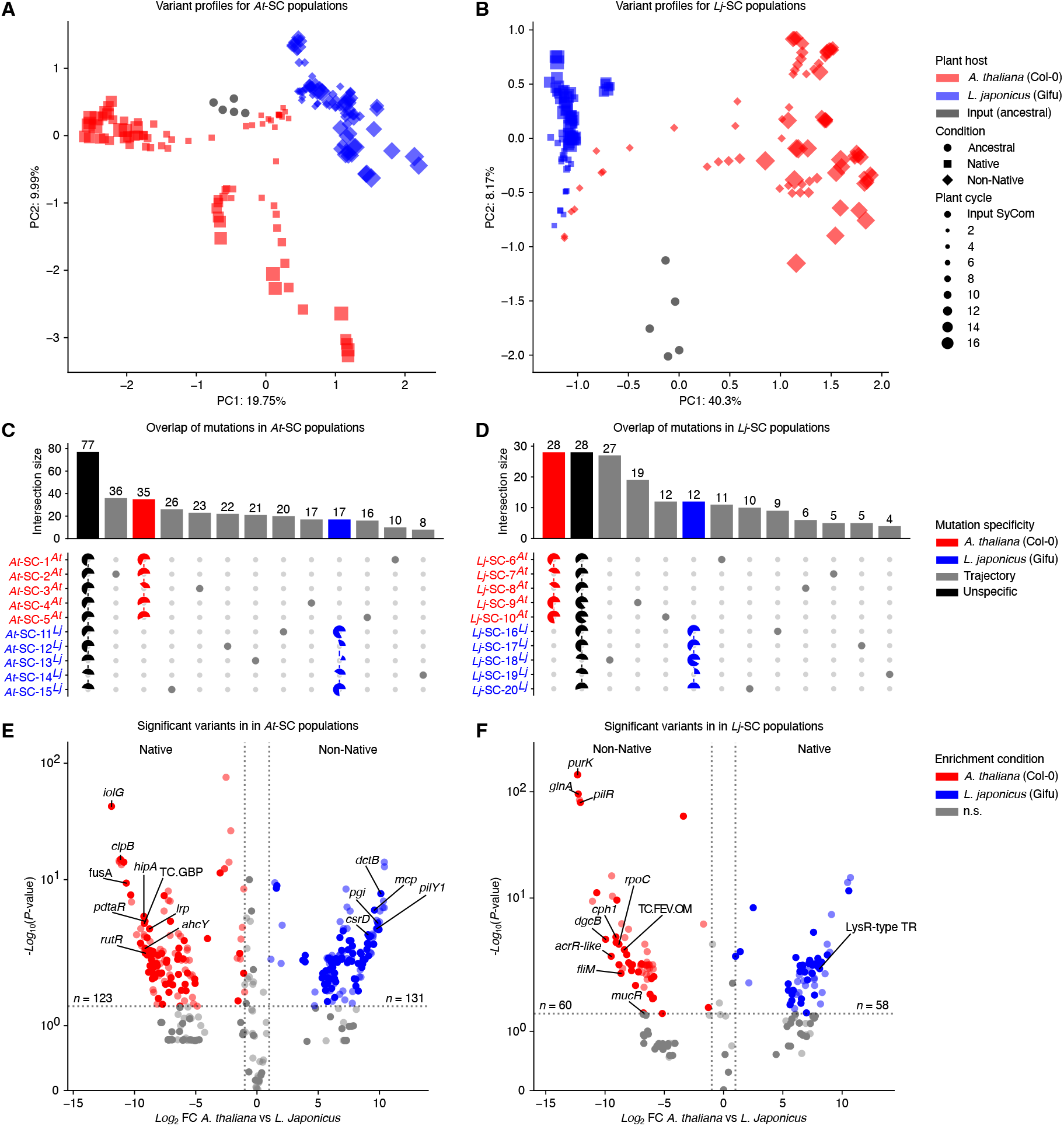
Host-specific variant profiles. Principal component analysis using a matrix of allele frequency values for SynComs *At*-SC (**A**; *n* = 163) and *Lj*-SC (**B**; *n* = 160). Different colours indicate the host plant the SynComs were evolving on during the evolution experiment. Point sizes represent the timepoint (*t*). (**C**-**D**) Upset plots showing the mutational overlap for the *At*-SC (**C**) and *Lj*-SC (**D**) SynComs on the level of open reading frames. Black bars and pie charts indicate mutated open reading frames that were found in both hosts. Red and blue elements represent ORFs that were hit by mutations exclusively during evolution either on *Arabidopsis* or *Lotus* roots. Grey elements were specific to a replicate population. Pie charts summarize the fraction of mutated ORFs that were found in a given replicate population. (**E**-**F**) Mutations significantly enriched on either *Arabidopsis* (red) or *Lotus* (blue) within *At*-SC (**E**) and *Lj*-SC (**F**). Enrichments were determined by fitting a generalized linear model per detected mutation (*n* = 543). *P*-values of enrichments were adjusted using false discovery rate (α = 0.05).

We hypothesized that such reproducible evolutionary outcomes could be explained by specific genetic variants appearing independently in multiple replicate lines. Supporting this hypothesis, we identified 197 bacterial genes that acquired high-confidence mutations in multiple populations compared to 307 genes with mutations in only one replicate line. Employing a binomial statistical test, we identified 259 genes (corresponding to 50.9% of all with detected mutations) showing significant patterns of non-random emergence across independent populations (**Table S2**; **Methods**). While some of these mutated genes were identified in either SynCom independently of the host species (77 in *At*-SC; **Figure 3C**, and 28 in *Lj*-SC; **Figure 3D**), we also identified mutated genes found exclusively in multiple replicate bacterial populations growing on their non-native host (17 for *At*-SC growing on *L. japonicus* roots; **Figure 3C**, and 28 for *Lj*-SC growing on *A. thaliana*; **Figure 3D**). These findings further support the hypothesis that repeated exposure to a new host species induces reproducible genetic changes in independently evolving bacterial communities.

Next, we performed a generalized linear model (GLM) analysis aimed at identifying genetic variants significantly enriched in either SynCom evolving on *Arabidopsis* or *Lotus* roots. We found 372 out of 543 (or 68.51%) variants that were enriched in bacterial populations associated with one host, indicating differential selection pressures operating on these communities (**Figure 3E-3F**; **Table S3**). This is consistent with the observed differences in fixation probabilities (**Figure 2J**), and the clear host-driven clustering patterns observed in the PCA plots (**Figure 3A-3B**). Collectively, these results suggest intensified selective pressures and greater adaptive potential when bacterial communities encounter a new host species.

To better understand the functional relevance of host species-enriched variants, we analysed their distribution across different gene categories. To specifically identify mutations that are adaptive in associations with a new host plant, we focused on mutations that emerged in non-natively evolving bacterial populations. A notable proportion of these variants affected genes encoding transcriptional regulators (e.g., LysR-type transcriptional regulators), and two-component systems (e.g., *pilR*, *dctB*). Such mutations have been observed in other experimental evolution systems, where they are thought to mediate rapid adaptation through modulation of transcriptional outputs, rather than innovation by acquisition of new functionality^17,18,24,25,42,46–51^. Besides mutations in genes related to transcriptional regulation, we also found a significant number of host species-enriched variants in genes related to motility and pilus assembly, and thus potentially relevant for root colonization (e.g., *fliM*, *fliE*, *fliF*, *pilN*), or involved in core metabolic processes in bacteria (e.g., *glnA*, *purK*, *bglB*, *icd*), which could play a role in adaptation to the distinct root exudation profile of a new host. Alternatively, these mutations could modulate bacterial cell surface structures and consequently, act as camouflage against the immune system of the new host. Collectively, these data support the hypothesis that repeated exposure of SynComs to a new plant species results in reproducible patterns of genomic adaptation.

### Increased fitness during colonization of a novel host species after experimental evolution

Next, we sought to experimentally validate whether the genomic changes observed during our evolution experiment translated into increased bacterial fitness during root colonization of a new host species. To achieve this, we conducted a series of competition experiments at multiple timepoints, assessing the capacity of evolved populations to compete for colonization on both plant hosts (**Methods**).

In these experiments, we co-inoculated axenic *Arabidopsis* and *Lotus* plants with independently evolving populations. Specifically, we matched populations evolving on their non-native host against each other (*At*-SC*^Lj^*vs. *Lj*-SC*^At^*), forcing them to compete for colonization of either plant species. As controls, we performed parallel competition experiments using populations evolving on their native host (*At*-SC*^At^* vs. *Lj*-SC*^Lj^*). After 5 weeks of plant growth, we assessed community composition through amplicon sequencing and estimated host preference as a proxy for bacterial fitness (**Methods**).

For the *Lj*-derived SynComs evolving on their non-native host (*Lj*-SC*^At^*), we observed a significant increase of 13.9% in bacterial fitness during colonization of *A. thaliana* roots by the end of the experiment (generation 16) compared to the start (generation 1; **Figure 4A**). A similar pattern was observed for the *At*-derived SynComs evolving on *L. japonicus* (*At*-SC*^Lj^*), which experienced an increase in fitness throughout the experiment of 27.5% (**Figure 4A**). This improvement in fitness emerged gradually and became statistically significant from generation 9 onwards, remaining stable until generation 16 (**Figure S5**). In contrast, for populations from both SynComs evolving in their native host (*At*-SC*^At^* vs. *Lj*-SC*^Lj^*) we observed no significant increase in fitness during colonization of the non-native host (**Figure 4B**), as expected given their lack of exposure to the new plant species.

**Figure 4.**
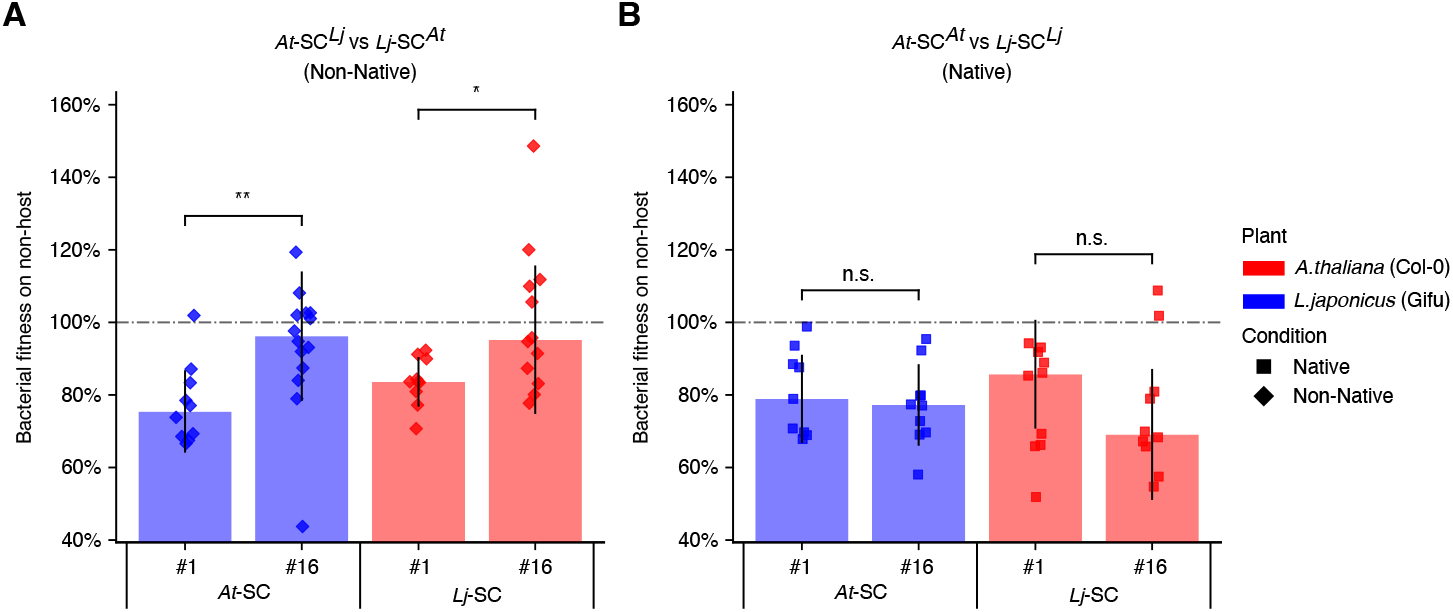
Bacterial fitness changes during experimental evolution. (**A**) Fitness changes of non-natively evolving *At*-SC*^Lj^*, and *Lj*-SC*^At^* (*n* = 22-24) at the beginning of the experiment (*t* = 1) and the end (*t* = 16) of the evolution experiment. Comparison was performed with a Mann-Whitney *U* test, and *P*-values were adjusted by calculating the false discovery rate (α = 0.05). (**B**) Fitness changes of natively evolving *At*-SC*^At^*, and *Lj*-SC*^Lj^*(*n* = 19-20). Data are represented as mean ± SD.

These results demonstrate that repeated exposure to a novel host species for 16 plant generations (∼20 months) is sufficient to induce measurable increases in bacterial fitness. Consequently, colonization efficiencies exhibited by evolved SynComs reach, or even surpass, those observed in native, host-adapted communities. Importantly, this phenotypic outcome aligns with the reproducible genomic signatures identified in our sequencing analyses (**Figure 3**), providing strong evidence that these genetic changes underpin enhanced fitness on novel hosts. Collectively, our results illustrate the rapid and reproducible adaptive evolution of synthetic bacterial communities to new host environments, driven by host-specific selective pressures.

### Identification and validation of adaptive mutations in evolved bacterial isolates

To independently verify our findings and identify genetic determinants underlying the observed changes in fitness, we isolated individual bacterial strains throughout the experiment (**Figure 1A**, **Figure 5A-5B**, **Figure S5A**, and **Figure S6**). We focused on isolates derived from the final plant generation (16), from which we validated and characterized a collection of 32 evolved clonal bacterial isolates. These isolates were subjected to whole-genome sequencing using long-read PacBio technology.

**Figure 5.**
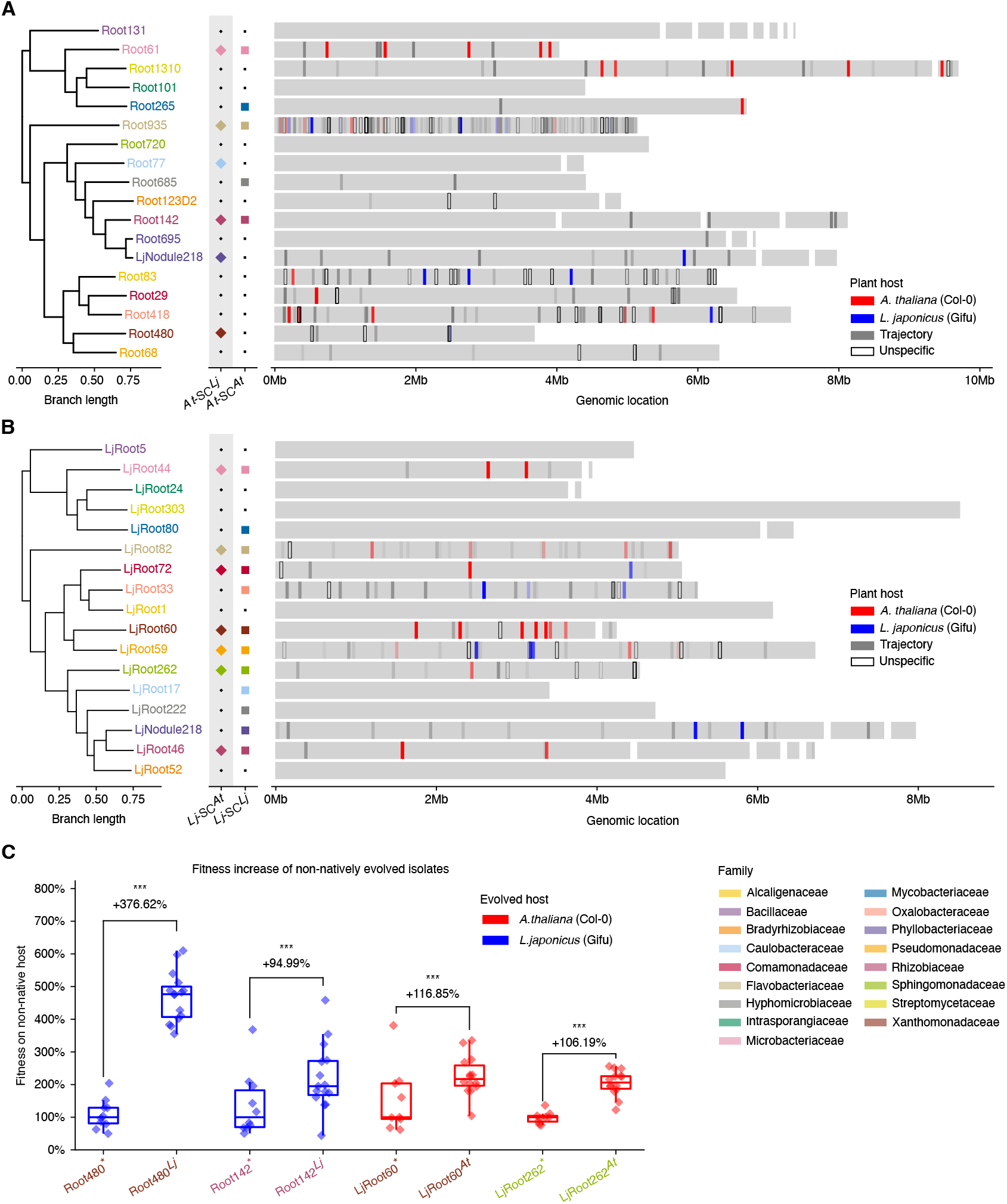
Genetic landscape and phenotype of evolved isolates. (**A**-**B**) Genomic context of detected mutations in the genomes of SynComs *At*-SC (**A**) and *Lj*-SC (**B**). Phylogenetic trees were inferred from all open reading frames. Coloured squares indicate strains for which an evolved isolate was obtained. Colouring of strain names and markers is according to taxonomy. Grey bars indicate the genomic structure of the respective strain. Red, blue, grey and transparent boxes mark candidate mutations that were determined from population level sequencing during the evolution experiment. (**C**) Bacterial fitness during root colonization of four evolved and reisolated strains that showed significantly increased competitiveness under non-native conditions (*n* = 9-15). Comparisons were performed between the ancestral and evolved strain using a one-sided Mann-Whitney *U* test, and *P*-values were adjusted by calculating the false discovery rate (α = 0.05). Data are represented as median with interquartile range.

Comparative analyses between the PacBio genome assemblies and earlier shotgun-sequencing data revealed that the majority (90.48%) of fixed variants identified through population-level sequencing were reliably found in the high-quality genome assemblies of the clonal isolates (**Figure 5A-5B**). We also analysed these genomes for large-scale structural rearrangements and signatures of horizontal gene transfer (HGT) events within our evolved bacterial strains. While structural rearrangements occurred sporadically (e.g., a large plasmid loss in strain LjRoot80; **Table S4**), we found no clear evidence of HGT events, suggesting limited genetic exchange within these synthetic communities during the evolution experiment.

To experimentally validate fitness changes observed at the population level (**Figure 4A-4B**), we reconstituted SynComs partially composed of evolved clonal isolates (mixed communities consisting of ancestral and evolved strains; **Figure 5A-5B**). We then performed competition experiments by matching non-natively (*At*-SC*^Lj^* and *Lj*-SC*^At^*) and natively evolved (*At*-SC*^At^* and *Lj*-SC*^Lj^*) reconstituted SynComs with their respective ancestral counterparts (denoted *At*-SC* and *Lj*-SC*) and inoculated them onto axenic *Arabidopsis* and *Lotus* roots. This experimental setup allowed us to assess changes in fitness of evolved clonal isolates compared to the ancestral strains in a community context. These experiments showed that SynComs evolved on a non-native host exhibited increased preference for that host compared to the ancestral SynComs (**Figure S7**), in line with previous results obtained from competition experiments of evolving populations (**Figure 4A**).

We detected four evolved strains with significant fitness increases during colonization of the non-native host (**Figure 5C**). Specifically, the strains Root480*^Lj^* (Xanthomonadaceae), and Root142*^Lj^* (Rhizobiaceae), originally derived from *A. thaliana* roots, increased their fitness in colonization of *L. japonicus* roots by 376.62% (*P* = 1.22×10^-4^) and 94.99% (*P* = 3.05×10^-4^), respectively. Similarly, strains LjRoot60*^At^* (Xanthomonadaceae), and LjRoot262*^At^* (Sphingomonadaceae) increased their fitness when colonizing their non-native host, *A. thaliana* by 116.85% (*P* = 1.52×10^-4^), and 106.19% (*P* = 1.53×10^-4^), respectively. Notably, these taxa were among the most abundant community members throughout the evolution experiment (**Figure 1C-1F**), suggesting that their high relative abundance and central ecological role may be related to their adaptive potential to new host environments.

Consistent with this increased fitness, PacBio sequencing confirmed the presence of adaptive mutations in several key genes within the genomes of these evolved strains, which were also previously identified in our population-level analyses via shotgun sequencing (**Table S5**). For instance, Root480*^Lj^* carried two mutations in genes related to glycogen metabolism (i.e., glycogen synthase and glycosyltransferase), which may alter carbon and energy storage in this strain, leading to adaptation to a new host environment with higher nutrient availability. Another example was found in LjRoot262*^At^*, which carried multiple mutations in the *cysJ* gene, encoding the sulphite reductase enzyme. We speculate that this could be the result of metabolic streamlining of this strain, originally isolated from roots of *L. japonicus*, in response to *A. thaliana* root exudates, which are rich in sulphur-containing glucosinolates^52,53^ and provide an alternative sulphur source that could be exploited by this strain. Together, these experiments validate both the changes in fitness (**Figure 4**), and the reproducible genomic signatures of adaptation (**Figure 3**) observed during the evolution experiment. Collectively, our findings indicate that the observed genomic and phenotypic changes are driven primarily by adaptation to the new host species, rather than artifacts of adaptation to our reductionist experimental setup.

## Discussion

The evolutionary dynamics of host-associated microbiomes remain poorly understood, despite their potential to shape microbiome function and host specificity over ecologically relevant timescales. Here, we directly address this gap by passaging replicated synthetic bacterial communities through two divergent plant host species for over 16 plant generations. We demonstrate that complex, multispecies microbiota can undergo rapid and reproducible adaptation to novel hosts, resulting in measurable gains in colonization fitness and the fixation of host-specific genetic changes.

This work builds on the LTEE framework, which has been applied mostly to single-strain^21,24–26,47^ populations or low-complexity, free-living microbial communities^27–29,31,33,34^, by extending it to a taxonomically diverse, host-associated microbiota. Unlike classical LTEEs, where adaptation is often traced to individual clonal lineages in well-mixed environments, our approach captures how selection unfolds across multiple interacting taxa in the structured context of the plant root. In such environments, microbial fitness is shaped not only by the capacity to colonize the host, but also by competition with other community members for host-derived resources^12,54,55^. Despite this complexity, we observed a surprising degree of evolutionary convergence. Replicate communities exposed to a new host follow parallel genomic trajectories (**Figure 3A-3B**), contain overlapping mutation targets (**Figure 3C-3D**), and undergo consistent gains in host species-specific colonization fitness (**Figure 4**). Together, these findings demonstrate that host-imposed selection can drive rapid and predictable adaptation across diverse microbial taxa, even in the presence of interspecific competition.

The reproducibility of these outcomes suggests that selection imposed by the plant host constitutes a strong and predictable force in shaping microbiome evolution even over timescales accessible to experimentation. This is further supported by the observation that community members evolved on their non-native host accumulated mutations more rapidly and fixed a greater proportion of them compared to populations maintained on their native host (**Figure 2J**). Notably, these patterns were not accompanied by major ecological disruptions. Community composition remained diverse and stable over time, with most founding strains persisting through all 16 plant generations (**Figure 1C-1F**). This ecological stability, paired with sustained genetic change, demonstrates that evolutionary adaptation can proceed without drastic community shifts. One possible explanation is that the community acts as a stabilizing force during evolution, where interspecific competition constrains reductive evolution and prevents the loss of costly competitive functions, a pattern often observed in simpler or single-species contexts^17,56,57^. In our experimental setup, the reproducible fixation of host-specific mutations likely reflects a balance between improving colonization of a novel host species and maintaining competitiveness within the existing microbial community.

Our findings demonstrate that root-associated microbiomes can evolve over short time periods. Combined with the well-known phenomenon of plant-soil feedback, the process by which plants modify the biotic and abiotic properties of their surrounding soil^58^, this suggests that host-driven selection continuously shapes soil microbial communities in natural ecosystems. We speculate that in sites where multiple plant species co-occur, repeated host filtering facilitates the formation of highly diversified soil biomes through ongoing, species-specific selection. Consistent with this prediction, in a single naturalized field site with 49 co-occurring plant species that have diverged for up to 140 million years, assembly of their root microbiota from the same soil biome differed both in diversity and composition and was positively correlated with host phylogenetic relatedness^59^. In agricultural systems, the capacity of root microbiomes to evolve over relatively short periods in associations with a new plant species may likewise allow crop rotations or spatial crop combinations to leave distinctive marks of host selection in the soil biome^60^.

At the genomic level, adaptive evolution is manifested in the repeated fixation of mutations in genes belonging to specific functional categories, particularly genes involved in transcriptional regulation, two-component signalling systems, motility, and core metabolism (**Figure 3E-3F**). Many of these categories have also been implicated in previous experimental evolution studies^17,24,25,47^, suggesting that similar genetic routes to adaptation may be shared across systems. However, the recurrence of these mutations across multiple independent populations, and their enrichment under non-native host conditions, highlights the importance of the host species in shaping evolutionary outcomes on the genomic level. In particular, our data indicate that adaptation to a novel host involves fine-tuning of regulatory networks and streamlining of metabolic capabilities, rather than acquisition of entirely new functions, a pattern consistent with a model of adaptation through optimization rather than innovation^19^.

Our experimental design further allowed us to directly link genetic change to phenotypic outcome. Through a series of competition assays, we show that SynComs evolving on a new plant species consistently outperform both ancestral communities (**Figure 5C**), and those evolved on their native host as controls (**Figure 4**). In several cases, we could trace these gains to putative adaptive mutations in genes related to nutrient assimilation and stress tolerance. For example, mutations in glycogen metabolism and sulphur reduction pathways may reflect selective responses to host-specific root exudates. These strains were among the most abundant community members throughout the evolution experiment (**Figure 1C-1F**), suggesting a link between ecological success (i.e., high relative abundance) and adaptive potential. Such findings point to the capacity for rapid functional tuning of bacterial strains within a community context, with implications for engineering or selecting beneficial microbiota.

Beyond the specific context of plant-microbiota interactions, our study contributes to a broader understanding of how evolution operates in host-associated microbial ecosystems. Unlike many free-living microbial populations, host-associated microbial communities experience repeated population bottlenecks, spatially structured environments, and fluctuating nutrient conditions, all factors that may constrain or direct evolutionary trajectories. Our data show that under such conditions, microbial communities can still evolve rapidly and predictably, even in a community setting. These results have conceptual and practical implications. Conceptually, they suggest that plant microbiomes are not only ecologically dynamic, but also evolutionarily plastic, and that hosts may exert selective forces that shape their microbiota over short timescales. This opens avenues for studying their co-evolutionary dynamics and raises questions about the extent to which host specificity and microbiome function are evolutionarily linked. Practically, our findings inform strategies for microbiome-based interventions in agriculture. They imply that microbial inoculants may be improved through directed evolution approaches that harness host-imposed selection, rather than relying solely on static strain collections or functional screenings.

Together, our results provide experimental evidence that root microbiomes can evolve rapidly and reproducibly, leading to significant increases in fitness in response to host selection. Our work also provides a number of candidate genes and functions that putatively mediate adaptation to a new plant species, which will need to be functionally validated in future studies. More broadly, our experimental setup establishes a framework for studying the evolution of complex communities in association with a eukaryotic host, enabling systematic interrogation of the rules that govern microbiome evolution.

### Limitations of the study

While our work provides a controlled model for studying microbiome evolution, it also raises new challenges. For example, it remains difficult to disentangle the contributions of interspecies interactions, spatial structure, and host factors to the observed evolutionary outcomes. Similarly, although we identify candidate adaptive mutations, functional validation remains labour-intensive, particularly due to the requirement for single- and multi-gene deletion mutants, which must then be tested in a community context. Finally, our use of synthetic communities, while necessary for tractability, does not fully capture the complexity of natural microbiomes. For example, our initial SynComs include single representative strains of the most abundant bacterial families present in the root microbiota, and do not include species-level variation at the start of the experiment. This could explain why we do not observe clear signatures of HGT during the evolution experiment, given that the likelihood of exchange of genetic material is lower between bacteria from different taxonomic groups^61,62^. Furthermore, because community composition was not a variable treatment in this study, we cannot definitively determine the extent to which the observed evolutionary trajectories were dependent on the specific composition of our SynComs^27,63^. It remains possible that a different set of initial bacterial strains would have driven evolution toward different outcomes, and future studies utilizing varying community composition would be needed to clarify this point. In addition, some microbial taxa observed in natural microbiomes have not yet been cultured^35,37^ and were therefore excluded from our experiment. Future work should extend this framework to more complex communities and additional hosts, and include transcriptomic, metabolomic, and integrated analyses of host responses to evolved communities.

## Resource availability

### Lead contact

Further information and requests for resources should be directed to and will be fulfilled by the lead contact, Ruben Garrido-Oter.

### Materials availability

This study did not generate any new, unique reagents.

### Data and code availability

Raw sequencing reads have been deposited at the European Nucleotide Archive (ENA) under accession numbers PRJEB98833 (amplicon sequencing), PRJEB98838 (shotgun sequencing) and PRJEB98839 (PacBio sequencing). Scripts for data analysis and plotting are available on GitHub: https://github.com/bdn-227/exp_evo_rootmicrobiota.

## Acknowledgements

We acknowledge C. Duran for the assistance during the evolution experiment, G. Altay for assistance in reisolating individual evolved isolates and the members of the lab for fruitful discussions. We thank Paul Schulze-Lefert for providing feedback on the manuscript. This work was supported by the German Research Foundation under the German Excellence Strategy (EXC 2048/1, project 390686111) and the European Research Council (project PHYCOSPHERES).

## Author contributions

N.K., K.W. and R.G.-O. conceived the research. N.K. performed all the experiments and analyses. R.G.- O. and K.W. interpreted the results. R.G.-O. and N.K. wrote the manuscript.

## Declaration of generative AI and AI-assisted technologies in the writing process

During the preparation of this work the authors used Google’s Gemini in order to refine certain sentences. After using this service, the authors reviewed and edited the content as needed and take full responsibility for the content of the publication.

## Supplemental information

Figures S1–S8

Tables S1-S8

## Methods

### Bacterial material and growth conditions

Bacterial SynComs were reconstituted from previously published culture collections^35,37^. The two SynComs, *At*-SC and *Lj*-SC, were designed to maximize taxonomic coverage while ensuring that every member had a unique 16S rRNA gene sequence and could be differentiated using amplicon sequencing. Each SynCom comprised 17 strains, representing 17 distinct bacterial families, with the same families represented in both *At*-SC and *Lj*-SC. The design also preserved compositional similarity to SynComs used in a previous study where they were demonstrated to show robust host preference phenotypes^35^, thus, facilitating comparative analyses. Lastly, to closely reflect natural conditions, the strains were picked with the explicit goal of maximizing the taxonomic coverage at the family level. Bacteria were cultured in TY-medium (5 g l^-1^ tryptone, 3 g l^-1^ yeast extract, 10 mM CaCl_2_), either in liquid culture or on agar plates with 15 g l^-1^ Bacto agar (Difco). Cultures were incubated at 25°C; liquid cultures were shaken at 180 rpm. We also determined the maximum *in vitro* growth rate of each strain by adjusting cultures to an initial OD_600_ of 0.02, incubating them in TY-medium at 25°C for 72 h, and recording absorbance every 10 minutes using a plate reader (Tecan Infinite 200 PRO). Maximum growth rates were estimated using the Growthcurver R package^64^. The ancestral genomes of all SynCom members were sequenced using PacBio HiFi and served as references for all downstream analyses (16S amplicon, shotgun metagenome, and whole-genome sequencing). Exact SynCom compositions and strain-specific physiological parameters are summarized in **Table S1**.

### Plant material and growth conditions

As host plants, *Arabidopsis thaliana* ecotype Columbia-0 (Col-0) and *Lotus japonicus* ecotype Gifu B-129 were used. Plant material and seed sterilization followed established protocols^35^. Briefly, *Arabidopsis* seeds were surface-sterilized with 70% ethanol for 5 minutes, washed twice in 100% ethanol for 1 minute, rinsed five times with sterile Millipore-filtered water (ddH_2_O), and stratified in darkness at 4 ^∘^C. *Lotus* seeds were scarified with sandpaper, incubated in 1% bleach for 20 minutes, and rinsed five times with sterile water. The sterilized *Lotus* seeds were germinated on sterile Whatman paper moistened with sterile ddH_2_O in square Petri dishes under short-day photoperiods (10 h light at 21°C and 14 h dark at 21°C). FlowPots containing the plant seedlings were placed into boxes (SacO_2_ TP5000) and incubated in walk-in growth chambers (SNIJDERSLABS) under short-day photoperiods (10 h light at 21°C and 14 h dark at 21°C).

### Gnotobiotic plant experiments

All gnotobiotic plant experiments were conducted using FlowPots^38^. Five days prior to the start of the experiment, peat was mixed with vermiculite in a 2:1 ratio. The mixture was then moistened with 200 ml ddH_2_O per litre to ensure efficient sterilization during autoclaving. FlowPots were assembled by filling each pot with 65 ml of the soil mixture, with a nylon mesh placed at the bottom to prevent clogging of the system. The assembled FlowPots were autoclaved once daily until the start of the experiment, for a total of four rounds of autoclaving. At the start of the experiment, each FlowPot was flushed with 50 ml of sterilized ddH_2_O and 50 ml of 0.2× Murashige and Skoog (MS) medium (0.88 g l^-1^ MS salts) via vacuum infiltration using a QIAGEN manifold (QIAvac 24 Plus, Qiagen). Subsequently, either three *Arabidopsis* or *Lotus* seeds were planted per pot. All procedures were conducted in a clean bench to ensure sterility throughout. Axenic plants were grown for two weeks under short-day conditions. Next, FlowPots were inoculated with either 20 ml of sterile 0.2× MS medium (axenic control) or 20 ml of 0.2× MS medium supplemented with the SynCom (evolution condition) inoculum. Inoculations were performed under sterile conditions in a clean bench. The pots were maintained under short-day photoperiods (10 h light at 21°C and 14 h dark at 19°C) for an additional five weeks until harvest. After a total of seven weeks of growth, shoot fresh weight was recorded for each individual plant per FlowPot. For 16S rRNA gene amplicon sequencing, the roots of the three plants within each pot were pooled into a single representative sample. Consequently, the maximum number of datapoints is 960 for shoot fresh weight and 320 for community profiling, respectively.

### Artificial selection of root-derived SynComs

The evolution experiment was conducted from April 2022 until December 2023 and comprised a total of 16 plant cycles (*t*), with each replicate line consisting of two FlowPots, and each FlowPot containing three plant individuals, on average. At the start, bacterial strains were streaked on individual TY-agar plates 14 days prior to inoculation. Taxonomy was confirmed via 16S rRNA gene amplification followed by Sanger sequencing of the obtained amplicons (sequencing performed at Eurofins; primers 799F and 1192R; ^37^). Bacterial cultures were inoculated as liquid cultures (10 ml of sterile TY-medium with 10 mM CaCl_2_) seven days prior to the FlowPot inoculation and grown at 25 ^∘^C while shaking at 180 rpm for seven days. Before the FlowPot inoculation, each liquid culture was washed twice in 10 mM MgSO_4_ and the optical density was adjusted. For the final inocula, bacterial strains were diluted in 0.2× MS medium to reach a final OD_600_ of 0.02. For all subsequent cycles, extracted root-associated microbial communities containing living bacteria were used as inocula for the next plant cycle, ensuring a continuous selection regime on the bacterial populations. For the extraction of live root-microbial communities, plant roots were dissected at the end of each plant cycle using sterilized forceps, tweezers, and a brush. After careful removal of all soil particles, roots were stored in phosphate-buffered saline (pH = 7) until all samples were processed. Roots were then dried on sterile Whatman paper and transferred into 250 µl of sterilized 10 mM MgSO_4_. Root-associated bacteria were released by grinding the roots with sterilized pestles. The resulting root slurry was split into four portions: (a) 10 µl or 5 µl were used for limiting dilutions of *Arabidopsis*- or *Lotus*-derived communities, respectively, with the purpose of counting Colony Forming Units (CFUs) and for re-isolation of evolved isolates; (b) 70 µl were used for inoculation of the next plant cycle; (c) 70 µl from *Arabidopsis* or 35 µl from *Lotus* roots were used for competition experiments; (d) the remaining 100 µl were used for DNA extraction and subsequent amplicon or shotgun metagenome sequencing. For inoculation of the next plant cycle, 70 µl of root microbial community was diluted into 40 ml 0.2× MS medium, and inoculation of two FlowPots was performed via vacuum infiltration (QIAvac 24 Plus, Qiagen). All steps involved in the harvesting procedure were carried out under sterile conditions. The 100 µl DNA samples were treated with 10 µl (30 Kunitz units) RNase-free DNase I (QIAGEN) for 10 minutes at 37°C to remove free-floating plant DNA. Samples were then snap-frozen in liquid nitrogen and stored at -80°C until further processing. DNA extractions were performed using the QIAamp DNA Micro Kit (QIAGEN; Protocol: Isolation of Genomic DNA from Tissues). Purified DNA was eluted in 30 µl nuclease-free water and stored at - 20°C until further processing. DNA samples from the evolution experiment were subjected to amplicon sequencing (in-house MiSeq instrument) and shotgun sequencing (Novogene).

### Competition experiments

To assess changes in bacterial fitness, we performed competition experiments between evolving populations throughout the evolution experiment. Evolving replicate populations of *At*-SC and *Lj*-SC were combined at selected plant cycles (1, 3, 5, 7, 9, 11, 12, 13, 14, 15, and 16) to yield 34-member communities. For these experiments, we mixed the natively evolving *At*-SC (*At*-SC*^At^*) with the natively evolving *Lj*-SC (*Lj*-SC*^Lj^*), as well as the non-natively evolving *At*-SC (*At*-SC*^Lj^*) with the non-natively evolving *Lj*-SC (*Lj*-SC*^At^*). Pairings of replicate populations (i.e., which *At*-SC and *Lj*-SC were matched in a replicate of the competition experiment) were determined using a pseudo-random number generator in Python. During a pilot experiment, we observed that CFUs derived from *Lotus* root samples were approximately two-fold higher than those from *Arabidopsis* roots. To establish an equivalent starting point for the competition experiments, 70 µl of the *Arabidopsis*-derived root microbial community was mixed with 35 µl of the *Lotus*-derived community. Importantly, the host preference phenotype, and thus our fitness assessments of differential colonization across plant roots, remained robust against such variations in input ratios^35^. FlowPots for the competition experiments were assembled, inoculated, and SynCom-infiltrated on the same day as the harvest of the evolving populations. Harvesting was performed as previously described^35^. In brief, plant roots were dissected using sterile forceps, tweezers, and a brush while submerged in sterile ddH_2_O. Soil particles were carefully removed, and roots were blotted on sterile Whatman paper to remove excess liquid. Dissected roots were transferred into Lysing Matrix E tubes (MP Biomedicals), snap-frozen in liquid nitrogen, and stored at -80°C until further processing. DNA extraction was carried out using the FastDNA Spin Kit for Soil (MP Biomedicals) following the manufacturer’s instructions, and DNA was eluted in 50 µl nuclease-free water. Samples were stored at -20°C until further processing.

### Re-isolation of evolved isolates

For re-isolation of evolved SynCom members, limiting dilutions were performed as previously described^65^. Briefly, 10 µl of homogenized root slurry from *Arabidopsis* or 5 µl from *Lotus* were first diluted in 1.25 ml of 10 mM MgSO_4_. This was followed by further dilution in 50 ml of 0.5× TY-medium (supplemented with 10 mM CaCl_2_) to reach the final concentration. For the 16th plant cycle, re-isolation efforts were doubled. Dilution factors in the second step were adjusted as needed across the experiment to ensure appropriate final concentrations (summarized in **Table S6**). For culture collection establishment, 160 µl of 0.5× TY-medium containing evolved strains was distributed into three 96-well plates (six plates for the 16th cycle). Plates were sealed with micropore tape (3M) and incubated at room temperature for two weeks. This procedure yielded 60 plates from plant cycles 1, 3, 5, 7, 9, 11, 13, and 15, and 120 plates from cycle 16. After incubation, wells showing visible growth were identified by scanning the plates with a plate reader (Tecan Infinite 200 PRO) to obtain fluorescence spectra and optical density data. Next, 150 µl of 80% (vol/vol) sterilized glycerol was added to each well. Plates were sealed with an aluminium seal (ThermoFisher) and stored at -80°C until further processing. Fluorescence spectra were used to classify all re-isolated strains, enabling high-throughput screening without the need to sequence all plate × well combinations. First, we collected training data ranging from 144 to 216 replicates per strain, as well as 432 replicates for medium-only controls (**Table S7**), yielding a comprehensive dataset for strain-specific classification. Spectral data were processed using custom Python scripts. Briefly, raw spectra were filtered to retain values within the linear range of the measurement device. To account for plate-to-plate variation, we performed medium-only measurements for each plate; these medium-only spectra were used to normalize the fluorescence data in a plate-specific manner. Standardization was performed by applying a *z*-score transformation followed by a hyperbolic arcsine transformation. The resulting normalized spectra were then corrected using a Python reimplementation of the ComBat function from the sva package^66^ to account for day-to-day measurement differences. The adjusted spectra were renormalized using a *z*-score transformation. The final spectra were used to train a random forest classifier with 500 estimators (scikit-learn v1.5.2). The resulting classifier was therefore capable of distinguishing wells containing only medium from wells exhibiting bacterial growth, and of discriminating among different taxa. We then applied the classifier to the fluorescence spectra obtained from the collection of evolved and re-isolated bacterial isolates (**Figure S6**). From these results, we estimated the number of wells lacking bacterial growth. Accounting for the applied dilution factors (**Table S6**), we inferred absolute bacterial abundances (i.e., colony-forming units) of root-associated microbial communities at the population level (**Figure S1C**). For recovery of specific strains of interest from the evolution experiment, glycerol stock plates were retrieved from the -80°C freezer, and TY agar plates were spot-inoculated and streaked using the three-way dilution method. After one week of growth, single colonies were picked, and taxonomic identity was confirmed via 16S rRNA Sanger sequencing. Verified colonies were inoculated into 5 ml liquid TY-medium supplemented with CaCl_2_ and incubated for one week at 25°C with shaking at 180 rpm. Glycerol stocks were prepared by mixing 800 µl of 80% sterilized glycerol (vol/vol) with 800 µl of culture and stored at -80°C for long-term preservation. The resulting culture collection of evolved strains comprised approximately 23,000 CFUs across all plant cycles. From this collection, 309 strains were re-isolated, re-streaked, and validated via 16S rRNA sequencing. Of these high-quality isolates, 34 were used for reconstitution experiments (**Table S8**), and 32 were subjected to whole-genome sequencing using PacBio HiFi sequencing (Max Planck Genome Centre).

### Reconstitution experiments with evolved isolates

Gnotobiotic reconstitution experiments were performed to independently validate the results of the competition experiments conducted during the evolution experiment. Axenic plants were grown for two weeks in FlowPots and inoculated with 0.2× MS medium (control) or with 0.2× MS medium containing different SynComs (**Table S8**), as described previously. Mixed communities consisting of ancestral and evolved strains were prepared using the same protocol used for initiating the evolution experiment. The design of the evolved communities (**Table S8**) was largely determined by the availability of sequence-verified evolved isolates at the time of the experiment. Our intention was to perform competition experiments with SynComs composed exclusively of evolved strains; however, two practical constraints prevented complete recovery of all evolved members. First, bacterial abundances on plant roots follow strongly skewed, log-shifted abundance distributions, which makes it challenging to recover low-abundance isolates. Second, evolved isolates can only be obtained from strains that persisted until the final plant cycle. Thus, taxa that went extinct during the evolution experiment are necessarily excluded. As a consequence, the competition experiments using evolved isolates, particularly experiment 1 (exp_1; **Table S8**), were carried out with SynComs only partially consisting of evolved strains. For taxa where no evolved representative could be recovered, we substituted with the ancestral strain, yielding SynComs that were evolved but partially ancestral. This strategy was chosen to eliminate SynCom diversity as a confounding factor. Thus, the competition experiments comprised 19 of 34 (56%) evolved strains. When accounting for extinction events (i.e., excluding taxa that went extinct during the experiment), these proportions increase to 19 of 29 (66%), representing the majority of community members. Harvesting was performed five weeks post-inoculation and followed the same procedure as used for the competition experiments during the evolution experiment.

### 16S rRNA gene amplicon sequencing

At each plant cycle, we performed amplicon sequencing of the bacterial 16S rRNA gene in order to characterize community structures. DNA samples were first adjusted to 1 ng µl^-1^ in nuclease-free water (QIAGEN). The 16S rRNA v5-v7 region was then amplified using primers 799F and 1192R with the following parameters: 94°C/2 min, 94°C/30 s, 55°C/30 s, 72°C/30 s, 72°C/10 min for 25 cycles^7^. Following amplification, PCR products were treated with 1 µl Antarctic phosphatase, 1 µl Exonuclease I, and 2.44 µl Antarctic Phosphatase buffer (New England BioLabs GmbH, Frankfurt, Germany). Next, 3 µl of the digested PCR product was used as template for a second PCR to add sample-specific barcodes (94°C/2 min, 94°C/30 s, 55°C/30 s, 72°C/30 s, 72°C/10 min for 10 cycles). The correct size of the amplicons from both PCR steps was confirmed using gel electrophoresis. After the second PCR, all samples were pooled to generate the final library. Gel electrophoresis and subsequent gel extraction using QIAquick (QIAGEN) were performed to remove chloroplast-derived 16S rRNA amplicons. The final library was purified with AMPure XP (Beckman Coulter) and eluted in 30 µl nuclease-free water. DNA concentration was quantified using QuantiFluor (Promega), and sequencing was performed on the in-house Illumina MiSeq platform.

### Processing of 16S rRNA gene amplicon data

Raw sequencing reads obtained from the MiSeq platform were demultiplexed using QIIME2^67^ v2021.11.0. Forward and reverse reads were subsequently merged with FLASH^68^ v2.2.00. Sequences containing undetermined nucleotides (i.e., “N” bases) were identified with usearch^69^ v10.0.240 and removed from the dataset. Reference-based error correction of high-quality sequences was performed using Rbec^70^ v1.1.4 with a subsampling depth of 500 and a Phred-scaled quality threshold of 33. The resulting count table was corrected for 16S rRNA gene copy-number variation and filtered for a minimal depth of 1000 reads. Subsequent downstream analyses were conducted in Python. Counts were normalized by calculating strain-specific relative abundances (RA). Stacked area plots (**Figure 1C-1F**) were plotted using ggplot2 and data preprocessing was performed using the *get_Muller_df* function of the ggmuller^71^ R package. Multivariate analysis (i.e., the effect of host plant on community structure) were performed using the R package vegan^72^.

### Host preference and fitness indices

We used a modified version of the host preference index^35^ to assess bacterial fitness. Specifically, fitness at the SynCom level (*At*-SC and *Lj*-SC) was estimated from differences in relative abundances observed in the competition experiments during the evolution experiment. For each sample *i*, we first calculated the aggregated relative abundance, 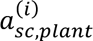, of all strains in SynCom *sc* on host *plant*. These values were then normalized by dividing by the median aggregated abundance of the same SynCom on its native host, *ã*_*sc*,native_, yielding the fitness score 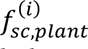 as defined in **Equation 1**. Here, *sc* ∈ {*At*-SC, *Lj*-SC} denotes the SynCom, *plant* ∈ {*Arabidopsis*, *Lotus*} indicates the host plant, *i* indexes individual samples, and “native” refers to the host on which the SynCom naturally evolved and were originally isolated from.

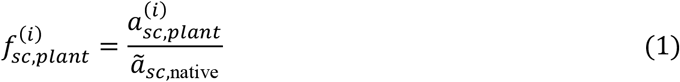

Differences in bacterial fitness were then investigated by comparing 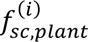 across different plant cycles (*t*) for a given condition. In particular, we focused on comparing fitness between the first and the last (sixteenth) plant cycle (plant cycle 1 vs. plant cycle 16; **Figure 4**).

For competition experiments with individually re-isolated evolved strains, fitness was calculated analogously to the SynCom-level fitness (**Equation 1**). Here, abundances of evolved isolates were normalized to the median abundance of their ancestral counterparts (plant cycle *t* = 0), yielding the fitness score 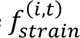 defined in **Equation 2**.

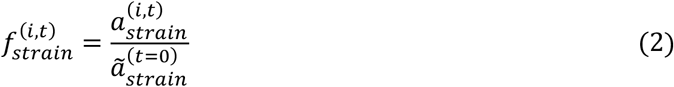

In this equation, 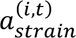 denotes the relative abundance of isolate *strain* (recovered at plant cycle *t*) in sample *i*, and 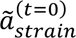 represents the median abundance of the ancestral, non-evolved strain. Here, *strain* ∈ {*At*-SC strains, *Lj*-SC strains}, *t* indicates the plant cycle from which the evolved strain was isolated, and *i* indexes individual samples. In practice, competition experiments with re-isolated strains were restricted to isolates from the final plant cycle (*t* = 16). To identify candidate evolved strains, we focused on non-natively evolved isolates that exhibited a statistically significant increase in fitness, i.e., for which 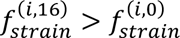 under the applied significance threshold, as determined by an adjusted *Mann-Whitney U* test.

### Processing of whole genome sequencing data

Raw HiFi sequencing reads were first filtered to remove the internal PacBio sequencing control by mapping the reads using blasr^73^ v1.3.1. Only unmapped reads were retained for downstream analyses. Assemblies were then performed using Hifiasm^74^ v0.25.0-r726 with default parameters and Flye^75^ v2.9.6-b1802 with coverage set to 70 and genome size set to 5 Mb. Both assembly graphs were visualized using Bandage^76^ v0.8.1 and manually inspected for structural issues such as bubbles. The quality of the assemblies was validated by re-aligning the filtered reads against the newly obtained assemblies using minimap2^77^ v2.28-r1209 (-ax map-hifi). Contigs with a mapping quality lower than 30 were considered assembly artifacts and removed. Circlator^78^ v1.5.5 was then used to scaffold circular, chromosome-level assemblies, and the remaining contigs were ordered by size in descending order. Assembly statistics were computed using assembly-stats^79^ v1.0.1, and the assembly with the highest *N50* was selected for downstream processing. The final assemblies were further evaluated for (a) contamination using CheckM2^80^ v1.0.1, (b) completeness using BUSCO^81^ v5.8.2, and (c) correct taxonomy using GTDB-Tk^82^ v2.1.1. Genome annotation was performed with Prokka^83^ v1.14.5 using an in-house curated KEGG-ontology database^84^. The same procedure was applied to evolved strains isolated from generation 16 of the evolution experiment. Mutations in evolved isolates were identified using two approaches: (a) assembly-based variant calling, where the assembly of the evolved genome was mapped against the assembly of the ancestral strain with Minimap2 (-x asm5) and calling variants using paftools.js^77^; and (b) read-based variant calling by mapping the HiFi reads against the ancestral assembly with Minimap2 (-ax map-hifi) and calling variants using DeepVariant^85^ v1.9.0 (- model_type=“PACBIO”). The resulting variant call sets were compared using custom Python scripts. Because the two sets were highly similar, only the read-based variant calls from DeepVariant were retained for downstream analysis, as this approach also allowed estimation of allele frequencies for identified variants. Lastly, we also investigated the occurrence of structural variation (e.g., horizontal gene transfer, duplications or larger insertions or deletions). Therefore, we aligned the reads obtained from evolved isolates against the ancestral genome using Minimap2. The resulting alignments were sorted by coordinates and used for structural variant calling using sniffles^86^ v2.6.3 with default parameters. Only structural variants with a homozygous genotype call (GT 1/1) were considered for downstream analysis. The majority of structural variants were deletions (12/13); however, we also detected one insertion. In order to identify the origin of the inserted DNA sequence, we used blast^87^ v2.14.1 to search the inserted DNA fragment in a concatenated reference containing all genome assemblies of the two SynComs. As the inserted sequence seemed to originate from the same genome, we concluded that these structural variants were not the consequence of horizontal gene transfer. Notably, the inserted sequence affected the coding sequence of a transposase, strongly suggesting an intra-genome rearrangement rather than horizontal gene transfer.

### Processing of shotgun sequencing data

Raw sequencing reads were subjected to quality control and host decontamination using KneadData v0.12.0 (https://huttenhower.sph.harvard.edu/kneaddata). Briefly, KneadData performed adapter trimming with Trimmomatic^88^ v0.39-2 (parameters: MINLEN:60, SLIDINGWINDOW:4:20), removal of tandem repeat sequences using TRF^89^ v4.09.1, host DNA removal with Bowtie2^90^ v2.5.3, and quality assessment with FastQC v0.12.1 (https://www.bioinformatics.babraham.ac.uk/projects/fastqc/). As host reference genomes, we used the *Arabidopsis* TAIR10 (Phytozome genome ID: 167; NCBI taxonomy ID: 3702) and *Lotus* Lj1.0v1 (Phytozome genome ID: 571; NCBI taxonomy ID: 34305) genome assemblies. Next, the decontaminated reads were mapped against a concatenated SynCom reference using Bowtie2. The resulting alignments were sorted with samtools^91^ sort v1.6, mate coordinates were added using samtools fixmate, and duplicate alignments were removed using samtools markdup. Potential mapping errors were corrected by performing probabilistic realignments with lofreq^92^ viterbi v2.1.5. Alignment and base-calling quality scores were then refined using lofreq indelqual (-dindel) and lofreq alnqual. The resulting alignments were used for all downstream analyses. Strain-specific sequencing depth and coverage were determined with samtools coverage. *In situ* growth rates were estimated using coPTR^40^ v1.1.6, which leverages uneven coverage distributions between the bacterial origin of replication and the terminus. First, decontaminated sequencing reads were mapped against the concatenated SynCom reference genomes using coptr map with default parameters. Growth rates were then calculated with coptr extract and coptr estimate with default parameters. Strain specific *in situ* growth rates were then parsed and analysed using custom Python scripts.

### Variant calling and analysis of shotgun sequencing data

Variant calling was performed using LoFreq, considering only mappings with a mapping quality ≥ 20 (-min-mq 20), using a *P*-value cutoff of 0.05 (-sig 0.05) and enabling indel calling (-call-indels). Potential multi-allelic variant calls were decomposed using vt^93^ decompose v0.5 (smart mode, -s), followed by normalization with vt normalize and removal of duplicate variants using vt uniq. Annotation of the candidate variants and effect predictions were performed with snpEff^94^ v4.3t (settings: -o gatk -noLog -noStats -no-downstream -no-upstream -no-utr -v -c). We further examined whether intergenic variants occurred in flanking regions of coding sequences to account for variants with potential effects on gene expression with a set of custom Python scripts. Finally, VCF files were parsed using custom Python scripts to obtain tab separated tabular files (tsv). Variant calling from population-level sequencing poses several challenges. In particular, the detectability of a variant is constrained by the sequencing depth of the respective genome relative to the total community depth. Consequently, variants from low-abundance taxa are easily underestimated when depth is limiting. To account for this effect, we performed variant-level rarefaction as follows. We generate *N* = 100 evenly spaced subsampling depths *dept*ℎ*_s_*

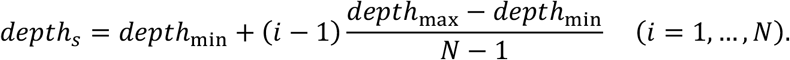

where *dept*ℎ_min_ corresponds to the number of reads for the shotgun sample with the lowest number of reads and *dept*ℎ_max_ to the highest observed number of reads. For sample *i* and subsampling depth *s* we compute the scaling factor *α_s_*_,*i*_

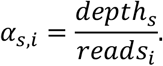

If *α_s_*_,*i*_ > 1 (i.e., the requested subsampling depth exceeds the raw reads of sample *i*), we skip that (*s*, *i*) pair (no up-sampling). For each variant *v* in sample *i* we denote the raw read support by

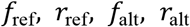

(forward / reverse reads supporting reference and alternative alleles). The *in silico* subsampled counts are then obtained by scaling:

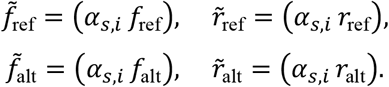

We then compute the subsampled depth (DP) and strand balance (SB):

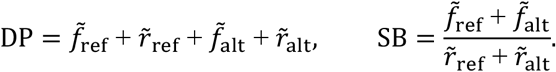

A variant passes the subsampled filter if

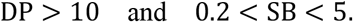

We repeat this procedure for all *N* subsampling depths in *dept*ℎ*_s_*, discarding variants for which *α_s_*_,*i*_ > 1. The pass/fail outcomes across subsampling depths provide a depth-aware measure of variant robustness. We visualized these results as rarefaction curves (**Figure S8**), plotting the number of detected variants per sample as a function of subsampling depth. Samples that reach a clear plateau (asymptote) at higher subsampling depths were considered to have sufficient coverage for robust variant calling. In contrast, samples that show an approximately linear increase in detected variants with depth are indicative of incomplete saturation and insufficient sequencing depth. Therefore, samples exhibiting no clear plateau were prioritized for additional sequencing to reach saturation. Additionally, we sought to independently validate the accuracy of the shotgun sequencing data. To this end, we compared the strain-specific relative abundances obtained from amplicon sequencing and shotgun sequencing. We observed a strong and highly significant correlation (*R^2^*= 0.844; *P* < 10^-308^) between the two methods, indicating that the metagenomic shotgun sequencing data are highly accurate.

Allele frequencies (*af_i_*) at a given genomic locus *i*, were calculated as the fraction of reads supporting a variant (*reads*_alt,*i*_) relative to the total number of reads covering that locus (*reads*_total,*i*_; **Equation 3**).

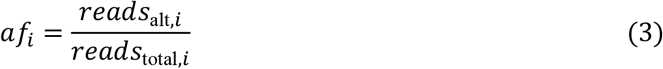

The variant call set was further filtered to include only variants supported by at least three mutant reads, with a strand bias between 0.2 and 5, a minimum Phred-scaled quality score of 30, a minimum total sequencing depth (i.e., the number of reads at the respective loci) of 10. In addition, variants had to be detected in at least four distinct timepoints within the same replicate population, and exhibit an allele frequency (AF) ≥ 0.1 in at least one plant cycle. To capture variants that emerged late in the experiment, the requirement for detection across plant cycles was relaxed: variants in generations 2 and 15 were required to be detected in at least three timepoints, whereas variants in generations 1 and 16 were required to be detected in at least two timepoints. Lastly, for downstream analysis we removed all variants that were already fixed in the starting inoculum to exclusively focus on variants that were being selected for during the experiment. For visualization of variant allele frequencies over time (**Figure 2A-2B**), missing values were imputed by linear interpolation where applicable; otherwise, values were set to zero for early plant cycles and carried forward from the last detected value for later plant cycles. Variant classes (**Figure 2G-2H**) obtained from snpEff and post-processed with custom Python scripts. To simplify analysis, related snpEff variant types were grouped into broader categories:

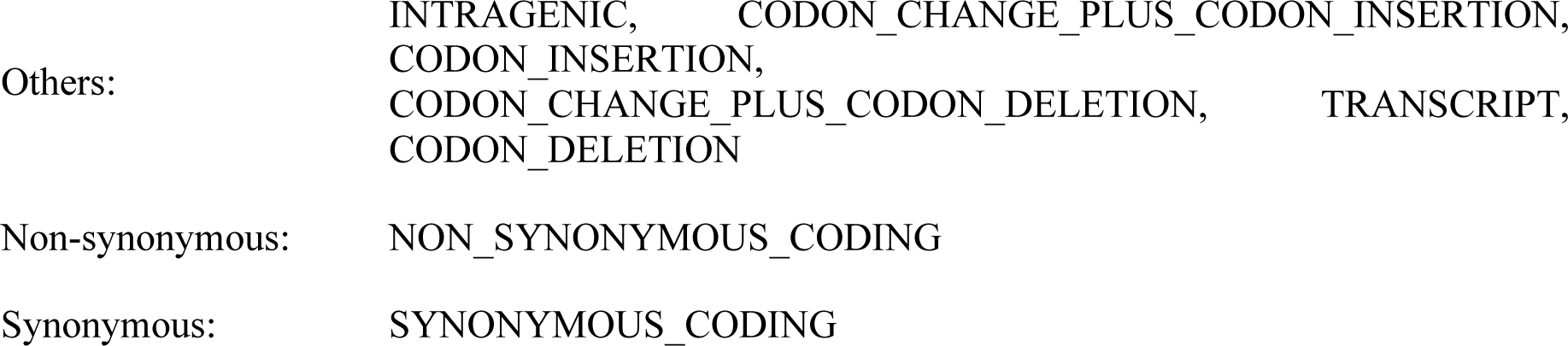
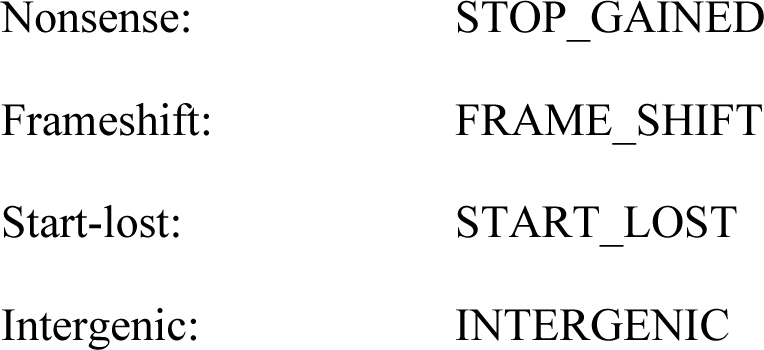

For each replicate population and timepoint *t*, we calculated the total derived allele frequency, *M(t)*, as the sum of all observed allele frequencies^22^. Fixation probabilities were calculated as the fraction of fixed variants (AF ≥ 0.95) relative to the total number of detected variants^22^. The resulting values were either aggregated at the strain level to obtain strain-specific fixation probabilities (**Figure 2I**) or at the condition level to obtain differential fixation probabilities for natively and non-natively evolved SynComs (**Figure 2J**). To estimate the proportion of variance in the variant call set explained by experimental conditions (host plant, SynCom, and plant cycle), redundancy analysis (RDA) was performed using scikit-bio^95^. Euclidean distances were calculated from the matrices of allele frequencies, followed by principal coordinate analysis (**Figure 3A-3B**). For the generation of the Upset plots (**Figure 3C-3D**), we first determined which replicate populations harboured candidate mutations in any given open reading frame (ORF). Next, we assessed whether a particular ORF was mutated exclusively in SynComs evolving on the same host and assigned a colour accordingly (red for SynComs evolving on *Arabidopsis* and blue for *Lotus*). ORFs that showed no pattern of host specificity were coloured black, while ORFs mutated in only a single replicate population were coloured grey. By convention, upset plots display all possible groups (in our case, all possible combinations of replicate populations with the same mutated ORFs). However, we chose to aggregate counts at the level of *Arabidopsis*-specific, *Lotus*-specific, line-specific, and unspecific ORFs. We summarized this information as pie charts (**Figure 3C-3D**). In this modified version of the Upset plot, each pie chart indicates the relative fraction of mutated ORFs occurring in the corresponding replicate population. To test for host-specific enrichment of candidate variants, we implemented generalized linear models (GLMs) using statsmodels^96^ v0.14.4. Each GLM was trained to predict allele frequency enrichment as a function of host plant. *P*-values were corrected for multiple testing across variants within each SynCom using false discovery rate (FDR) control. Data visualization was performed using Seaborn and Matplotlib using Python.

To assess non-random patterns of mutation occurrence, we applied a binomial test in a strain-specific and ORF-specific manner. Specifically, the probability *P*(*X* ≥ *k*) of observing *k* or more mutations in a given ORF was calculated using the **Equation 4**.

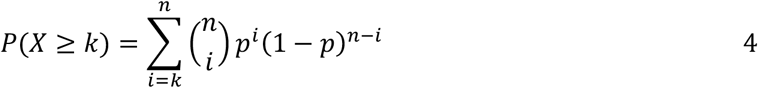

The total number of trials (*n*) was defined as the total number of mutations observed in the respective strain across all replicate populations. For each ORF, *k* (the number of successes) was defined as the number of replicates in which a mutation was detected within the respective ORF. The expected success probability, *p*, was calculated as the ratio of the added length (in base-pairs) of the ORF (*lengt*ℎ_ORF_) and its upstream region (*lengt*ℎ_upstream_) to the total genome length in nucleotides (*lengt*ℎ_genome_), as shown in **Equation 5**.

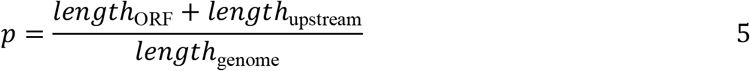

To reduce false positives arising from singleton events, only ORFs with *k* ≥ 2 were considered in the analysis. *P*-values, from the binomial test were corrected for multiple comparisons across ORFs using the false discovery rate (FDR) control.

### Data visualization

Plotting was, with the exception of **Figure 1C-1F**, performed using the Matplotlib and Seaborn ecosystem under Python v3.12.10. Phylogenetic trees (**Figure 5A-B**) were plotted using Biopython^97^ v1.74. As tree we used the species tree obtained from Orthofinder^98^ with default parameters v2.5.5 performed in the two SynComs (*At*-SC and *Lj*-SC). Boxplots display the median (horizontal line), with boxes spanning the interquartile range (Q1 - Q3) and whiskers extending to Q1 - 1.5× interquartile range (IQR) and Q3 + 1.5× IQR; individual datapoints are shown as circles or triangles with jitter. Schematic images (i.e., **Figure 1A-1B**; **Figure S5A**) were generated with Biorender. Final figures were assembled using Inkscape.

### Statistical information

For statistical inference, non-parametric tests were applied where appropriate. Two-group comparisons (including fitness changes during and after the evolution experiment, plant phenotypes, alpha diversity, and fixation probabilities) were evaluated using the Mann-Whitney *U*-test. Fitness indices between ancestral and evolved isolates (**Figure 5C**) were assessed using a one-sided test. The fixation probability of LjRoot33 and its hypermutator (**Figure 2I**) was evaluated via a one-sample *t*-test. Correlations (e.g., growth rates *in vitro* vs. *in situ*, growth rate vs. *M(t)*, relative abundance vs. *M(t)*, and CFUs vs. time) were quantified using Spearman’s *r*. Variant enrichments were determined using generalized linear models (GLMs), and non-randomness was evaluated via binomial tests. *P*-values were corrected for multiple testing using the Benjamini-Hochberg false discovery rate (FDR) procedure, with significance set at *α* = 0.05 unless stated otherwise.

## Supplemental information

### Figures

**Figure S1.**
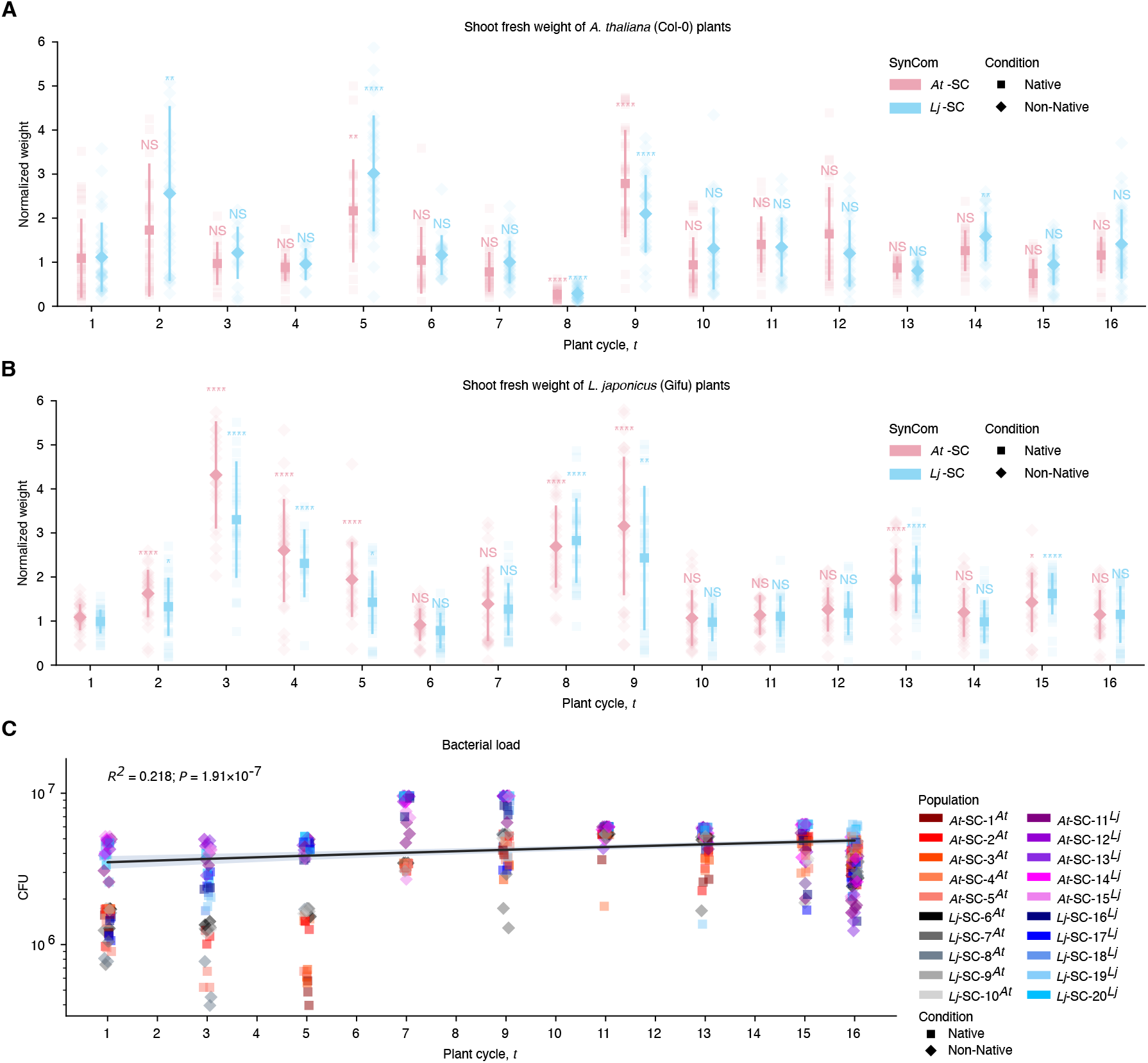
Plant fresh weight (**A**-**B**) and bacterial CFU counts over time (**C**). Plant weight (*n* = 875-897) was normalized to axenic controls. Statistical comparisons were made against the first plant cycle, with false discovery rate correction (*α* = 0.05). CFU values (*n* = 560) were estimated from reisolated strain cultures by adjusting the number of wells with visible growth for the dilution factor. Temporal trends were assessed using Spearman’s rank correlation between plant cycle and CFU count. Data are represented as mean ± SD.

**Figure S2.**
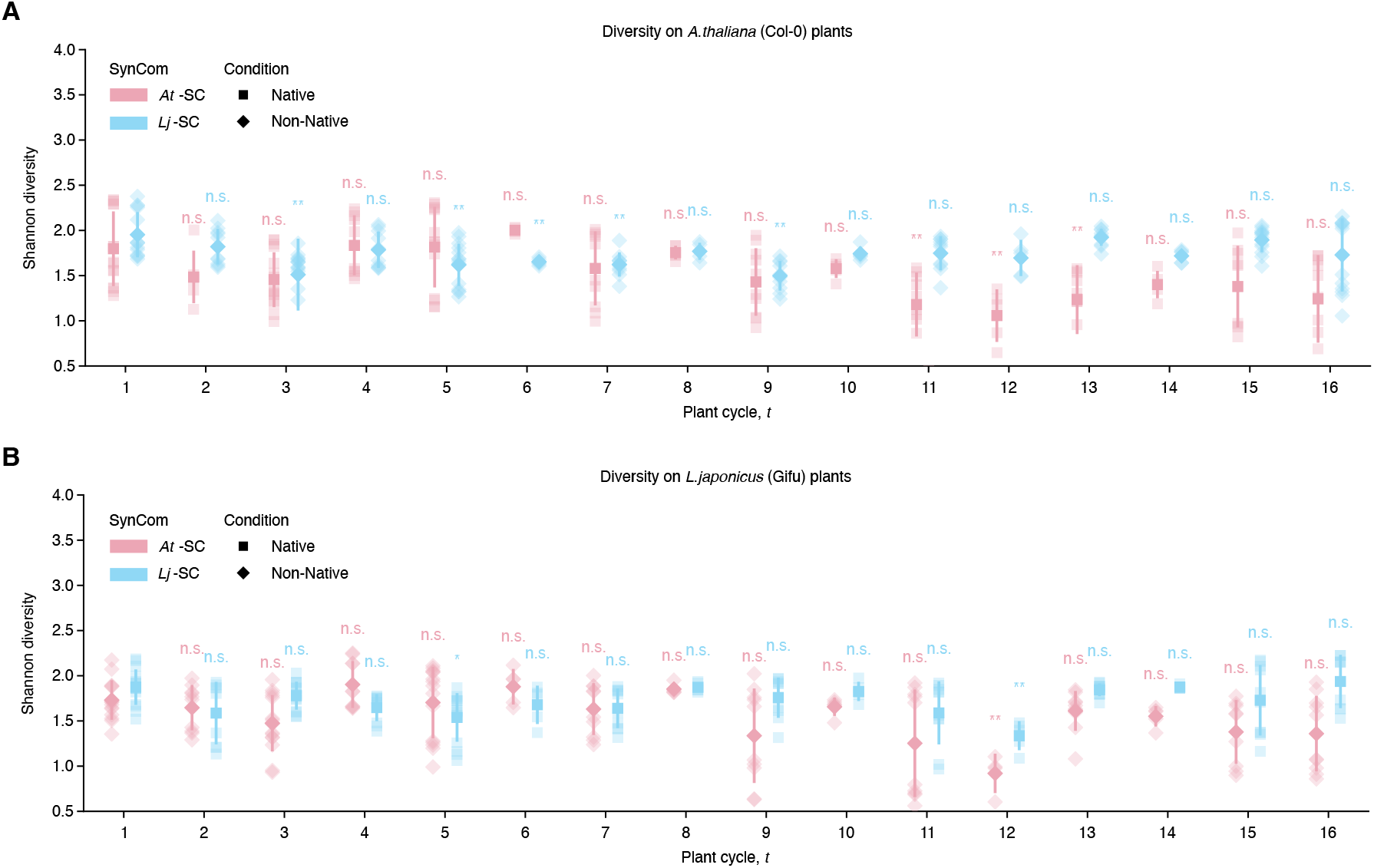
Bacterial α-diversity of evolving populations on *Arabidopsis* (**A**) and *Lotus* (**B**) obtained from amplicon data. Shannon entropy was calculated for SynComs evolving on both host plants (n = 294-299). Statistical comparisons were made against the first plant cycle, with P-values adjusted for multiple testing using false discovery rate control (α = 0.05). Data are represented as mean ± SD.

**Figure S3.**
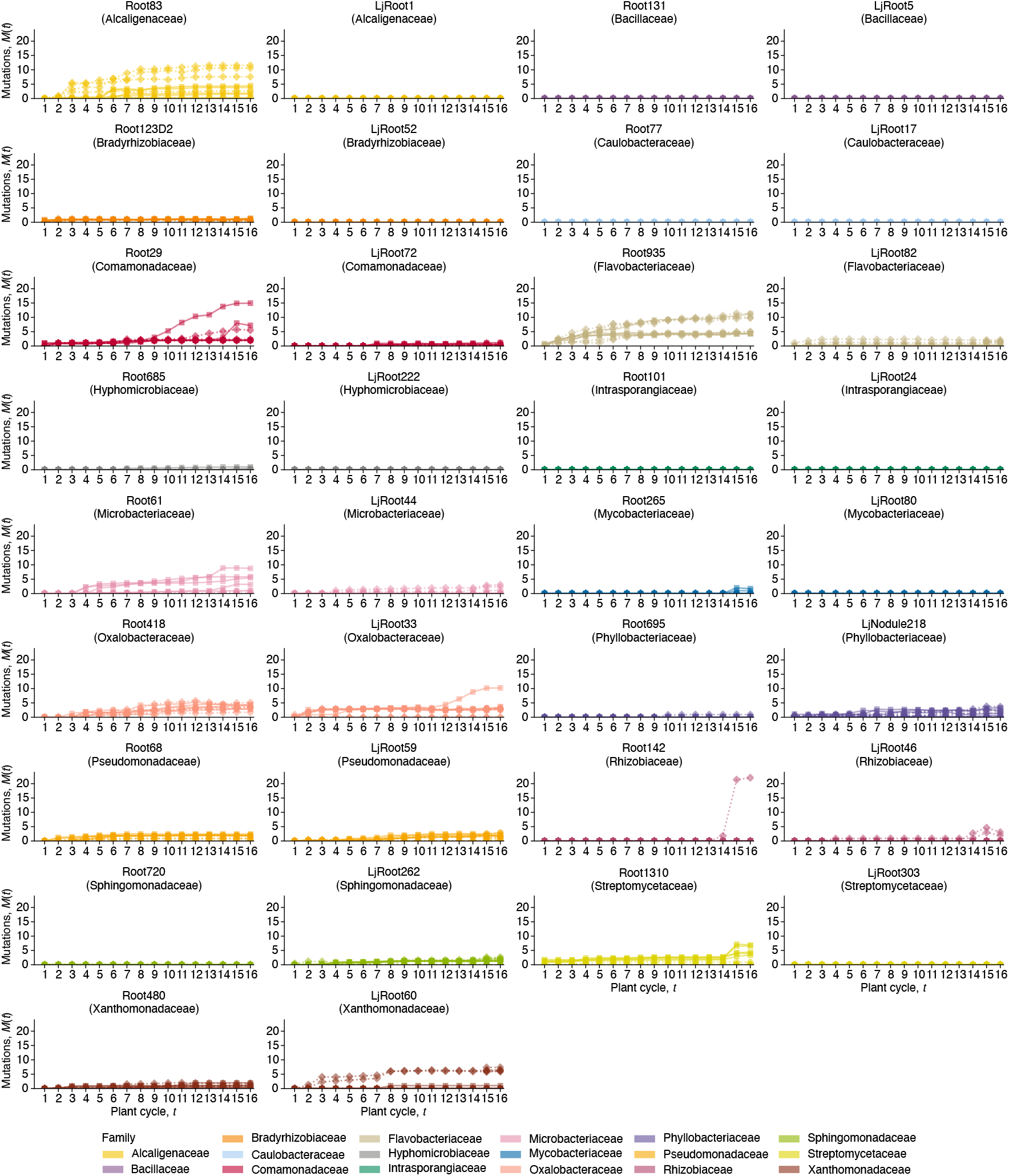
Strain-specific patterns of variant accumulation over time. Total derived allele frequencies *M(t)* are plotted against plant cycle for each strain. Lines represent the temporal trajectory of mutation accumulation within individual genomes.

**Figure S4.**
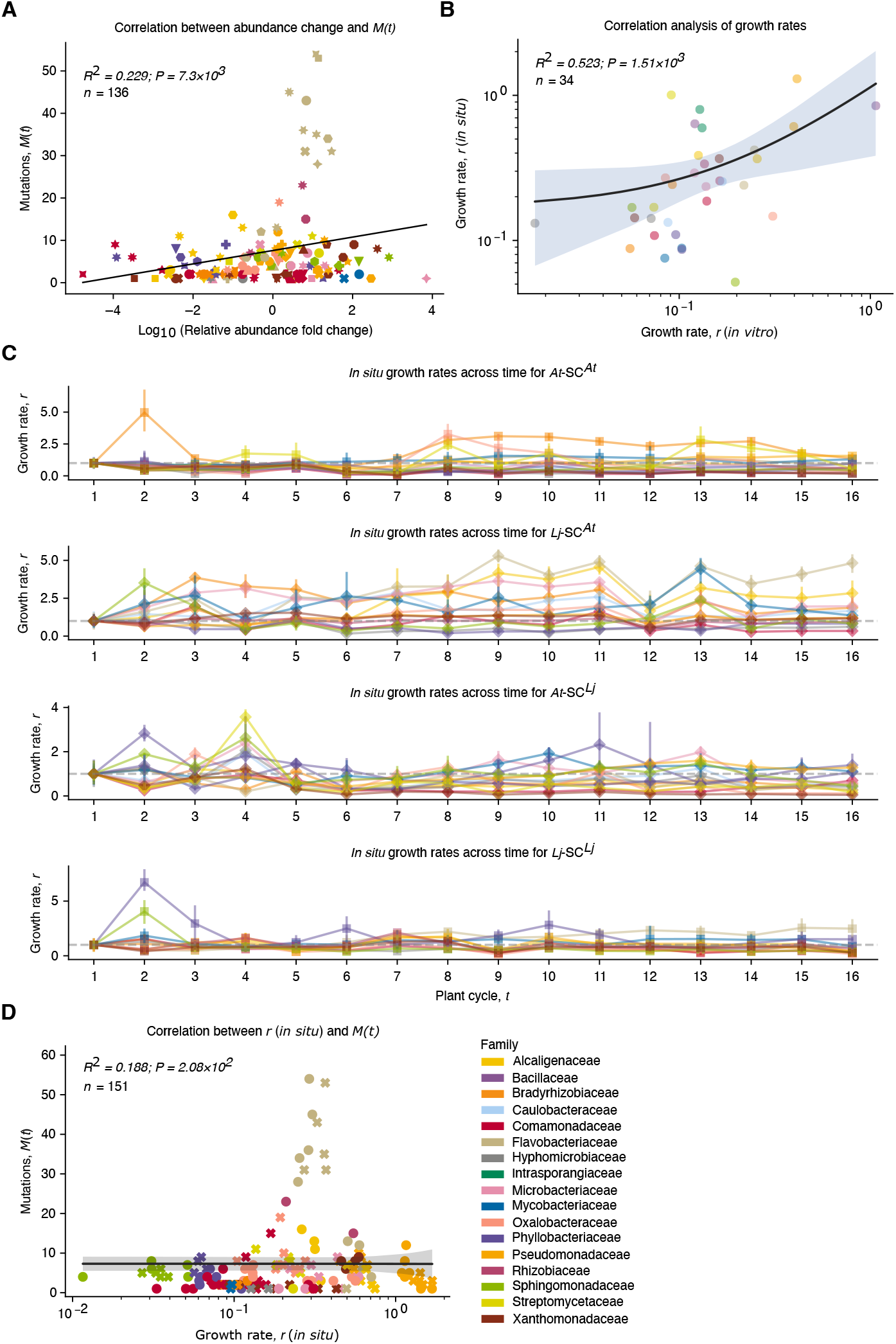
Bacterial growth rates during the evolution experiment. (**A**) Correlation between fold change in relative abundance and total derived allele frequency *M(t)* (*n* = 136). (**B**) Correlation between *in vitro* and *in situ* growth rates (*n* = 34). The black line represents the linear regression, while the shaded area indicates the 95% confidence interval. (**C**) *In situ* growth rates across plant cycles (*n* = 4508). Data are represented as mean ± SD. (**D**) Correlation between in situ growth rates and *M(t)* (*n* = 151). The black line represents the linear regression, while the shaded area indicates the 95% confidence interval.

**Figure S5.**
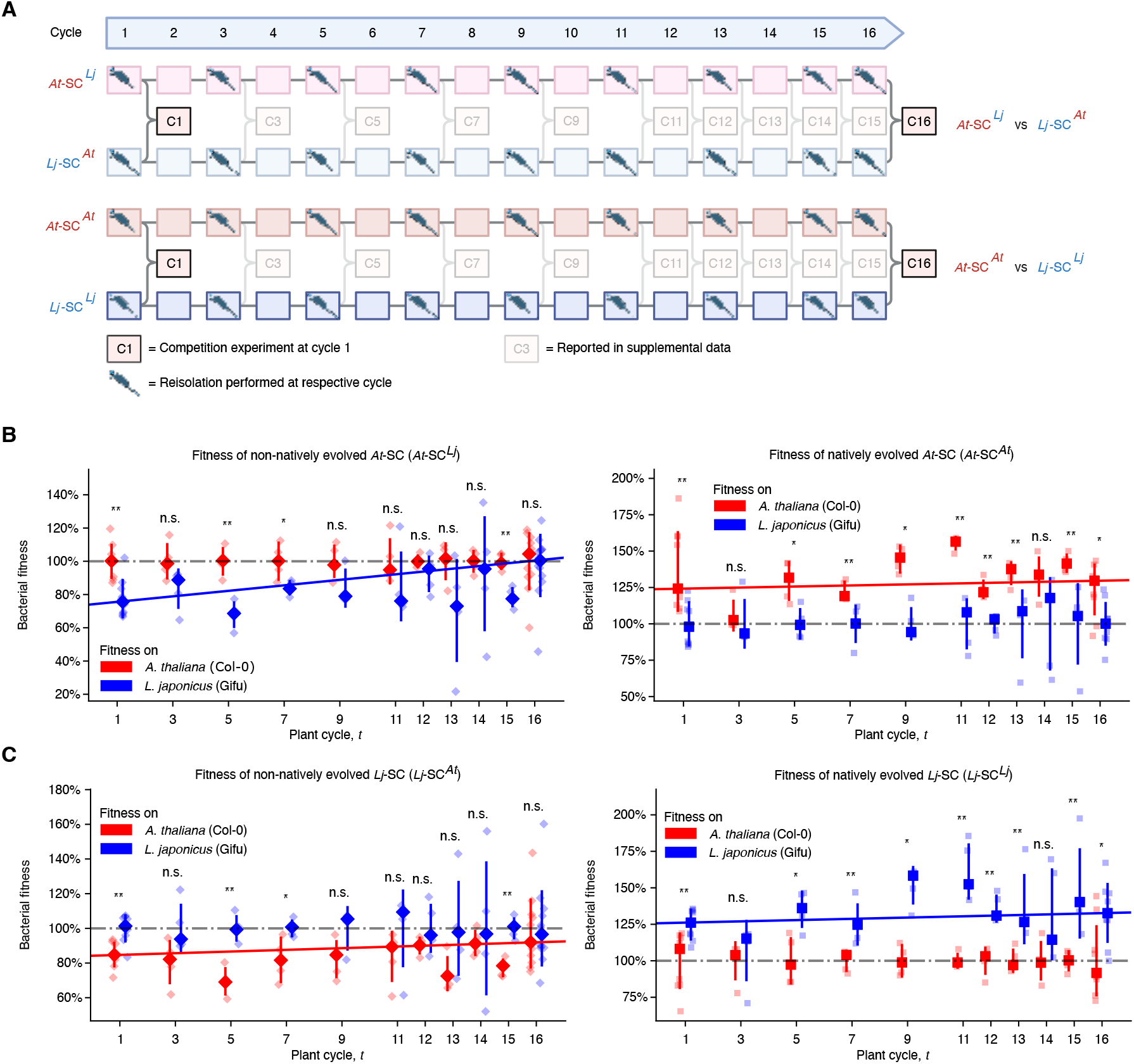
Changes in host preference of evolving populations. (**A**) Schematic of competition experiments indicating timepoints and mixed conditions. Pipette symbols mark collection of evolved isolates via limiting dilution. (**B**-**C**) Fitness trajectories of *At*-SC and *Lj*-SC evolving under native and non-native conditions (*n* = 128-134). Statistical comparisons assessed SynCom fitness on *Arabidopsis* versus *Lotus* roots (colours). Upper panels show fitness on non-native hosts, lower panels on native hosts. Coloured lines depict fitted trajectories of bacterial fitness on the experimental evolution host over time. Data are represented as mean ± SD.

**Figure S6.**
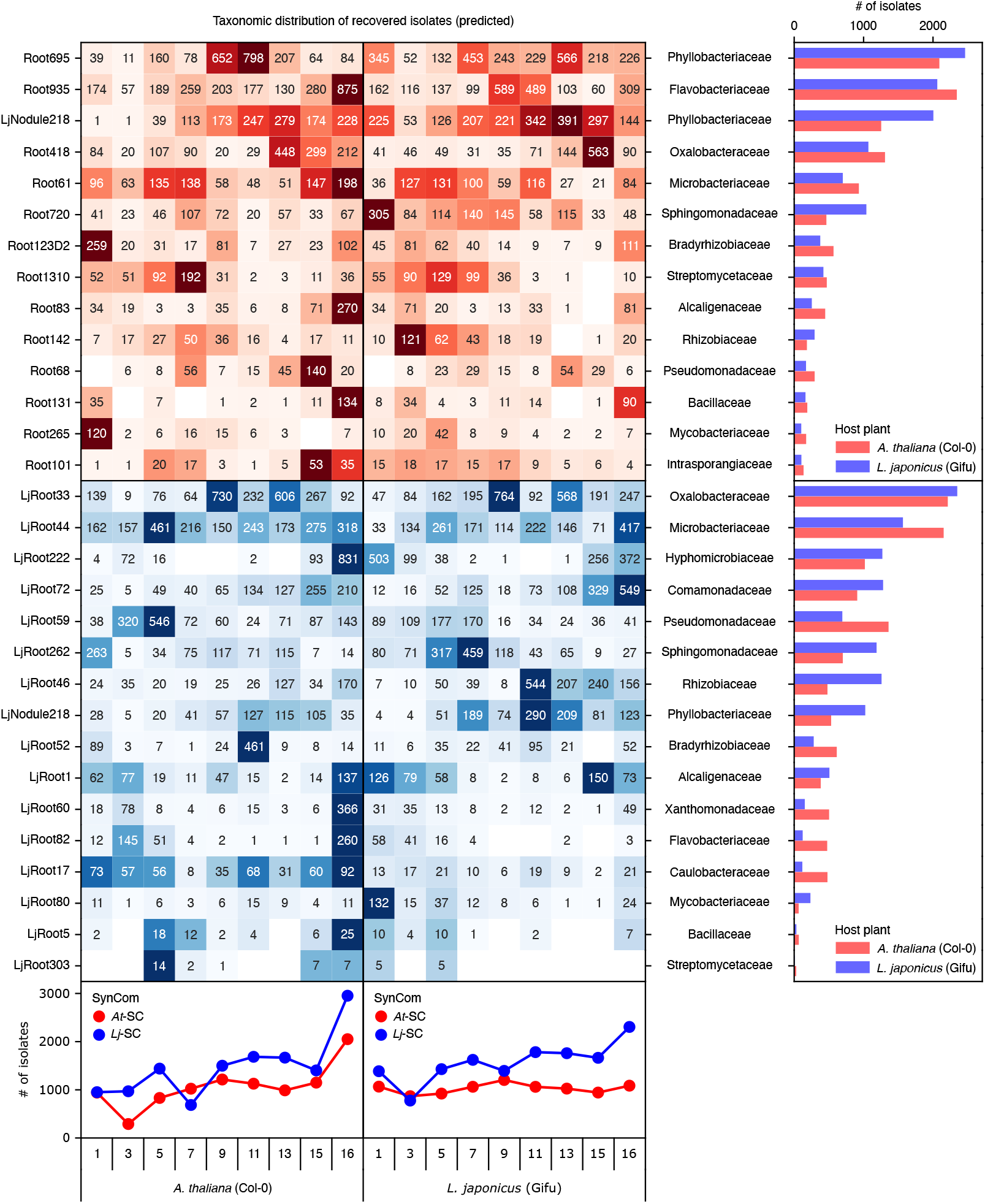
Taxonomic distribution of evolved and reisolated strains. Heatmap showing the predicted number of isolates per taxon at each timepoint. *At*-SC strains are red, *Lj*-SC strains are blue. Aggregated counts per SynCom and host are shown below the heatmap, and total counts per taxon across all timepoints are shown on the right.

**Figure S7.**
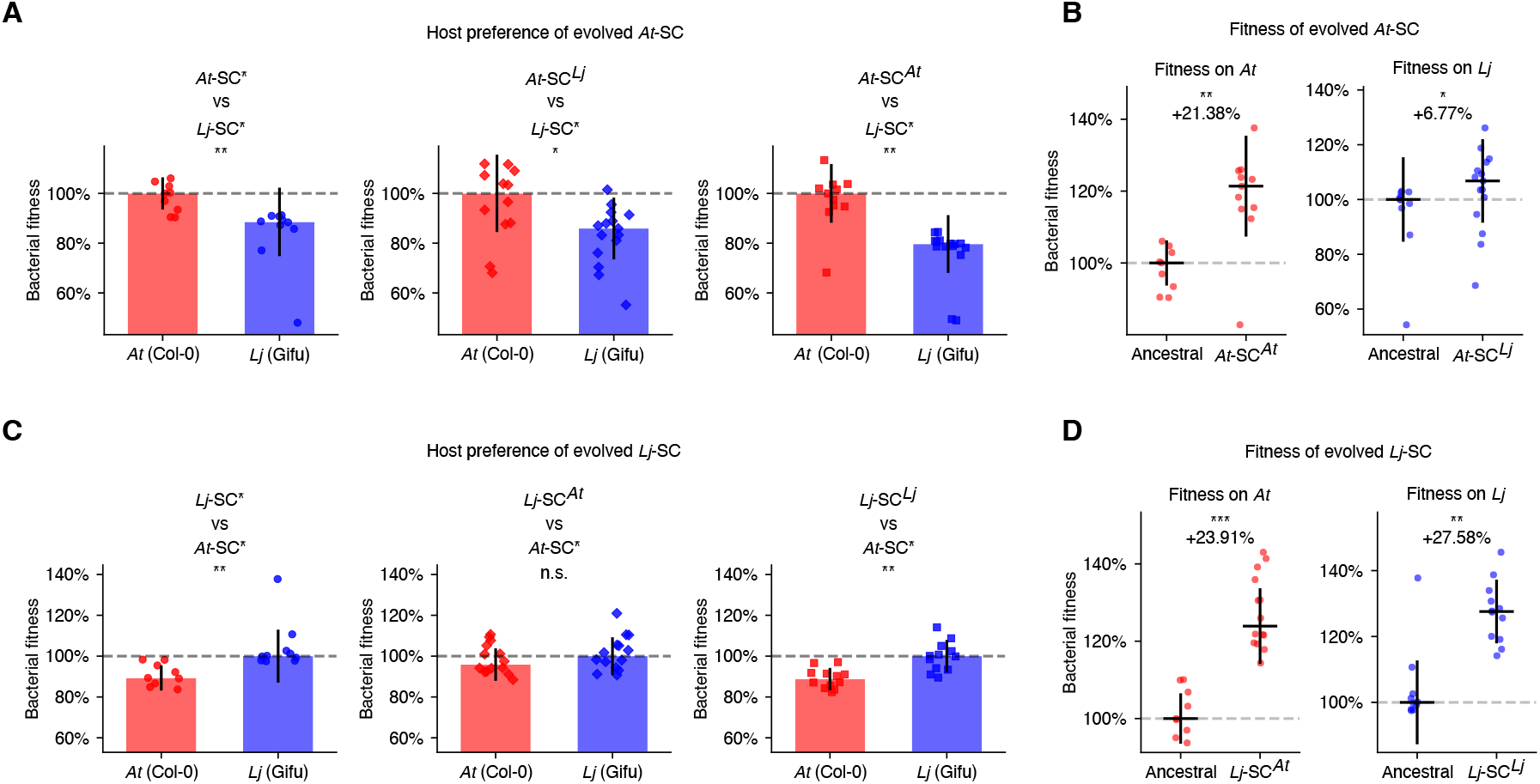
Fitness changes in SynComs containing evolved isolates. (**A** and **C**) Host preference of evolved isolates from *At*-SC and *Lj*-SC (*n* = 10-15). The first panel shows host preference of the ancestral strains; the second, competitions between non-natively evolved isolates and the ancestral counterpart of the other SynCom; and the third, competitions between natively evolved isolates and the corresponding ancestral SynCom. Fitness values were normalized to the native host. (**B** and **D**) Fitness of evolved *At*-SC and *Lj*-SC on both hosts, normalized to their ancestral fitness. Multiple testing correction was applied using false discovery rate control (*α* = 0.05). Data are represented as mean ± SD.

**Figure S8.**
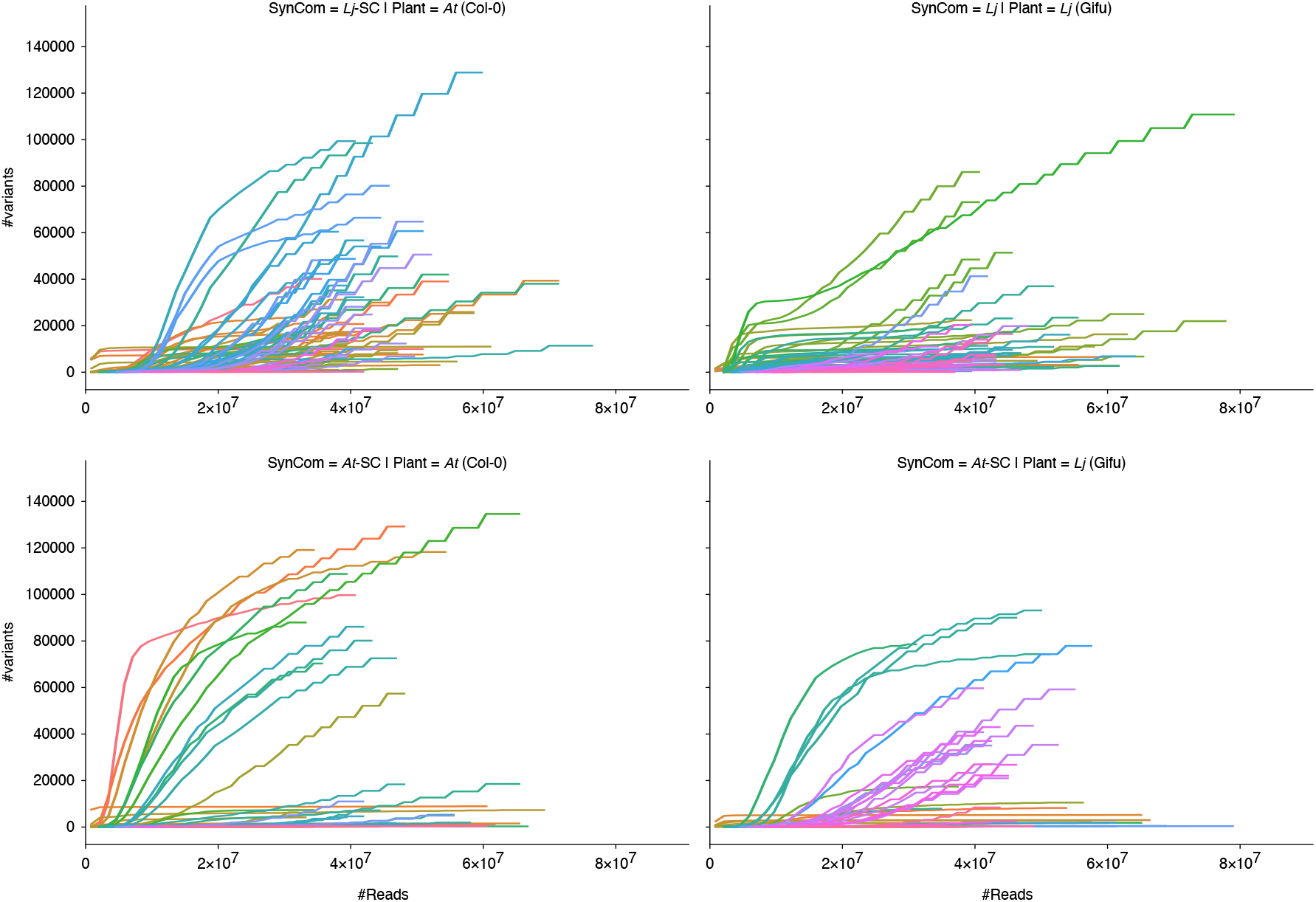
Rarefaction curves of variant calls. The number of detected variants (*y*-axis) is plotted against subsampling depth. Each line represents a shotgun sequencing sample. Analyses were performed for both SynComs (*At*-SC and *Lj*-SC) evolving on both hosts, resulting in four facets.

### Tables

**Table S1.**
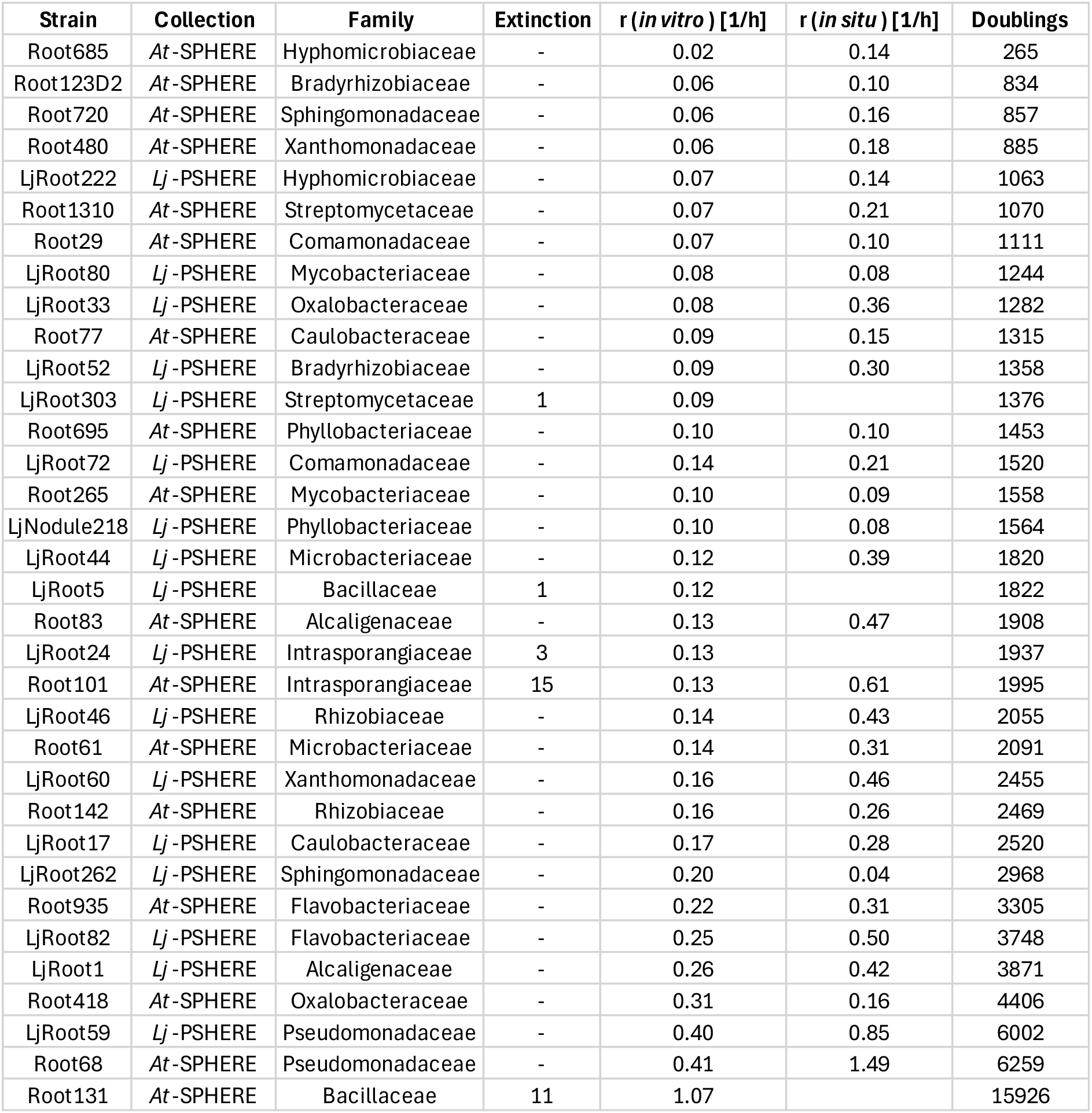
Summary of the SynComs used in this study. Strain name, native host, and taxonomic family are listed (*n* = 34). If extinction events occurred, the generation at which they occurred is indicated in the respective column. Strain-specific *in vitro* growth rates were inferred from real-time OD measurements of clonal strains in microtiter plates. *In situ* growth rates were calculated from uneven coverage distributions observed in population-level shotgun sequencing data. From these rates, the estimated number of bacterial doublings throughout the evolution experiment were calculated.

**Table S2.**
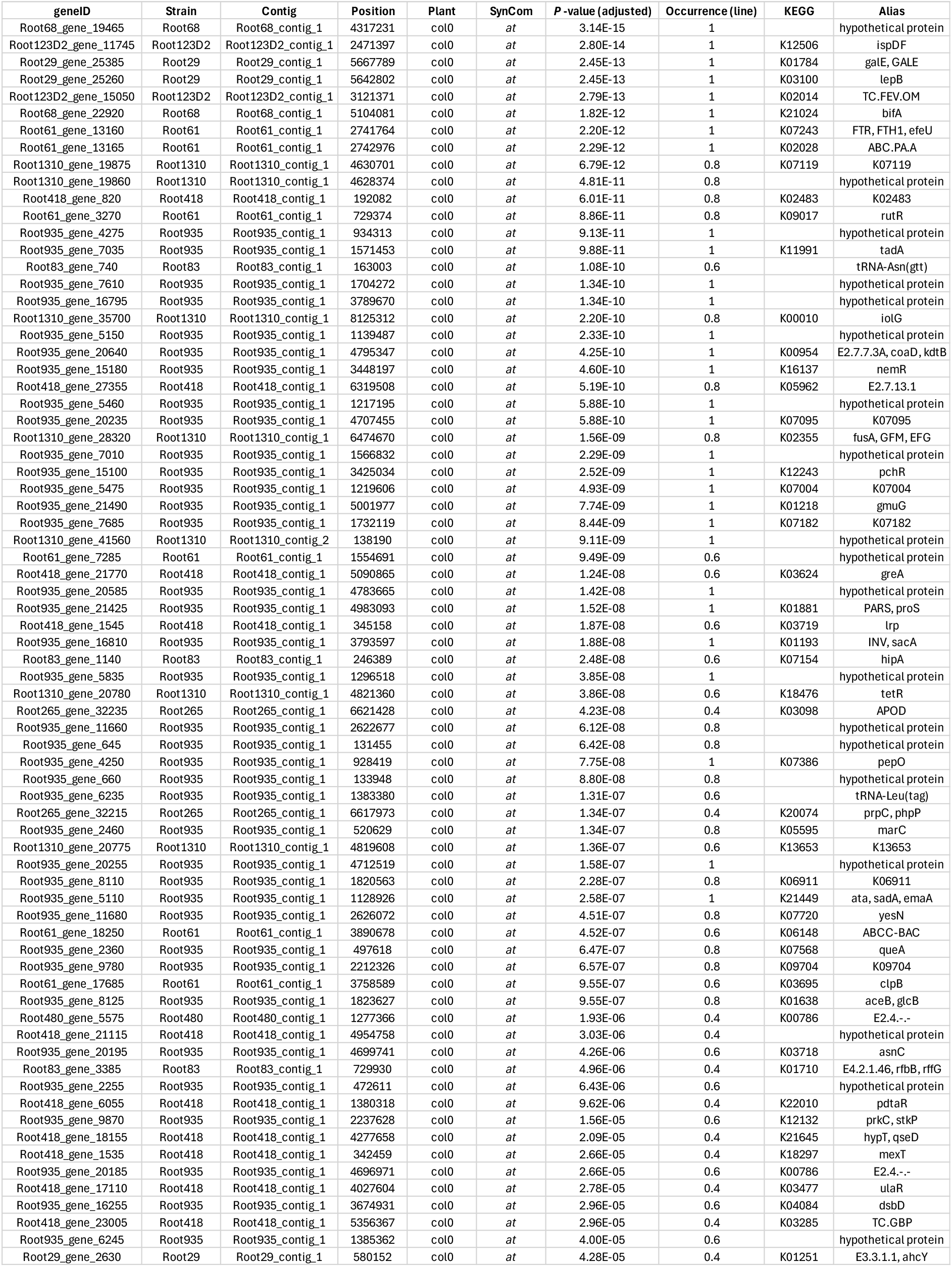
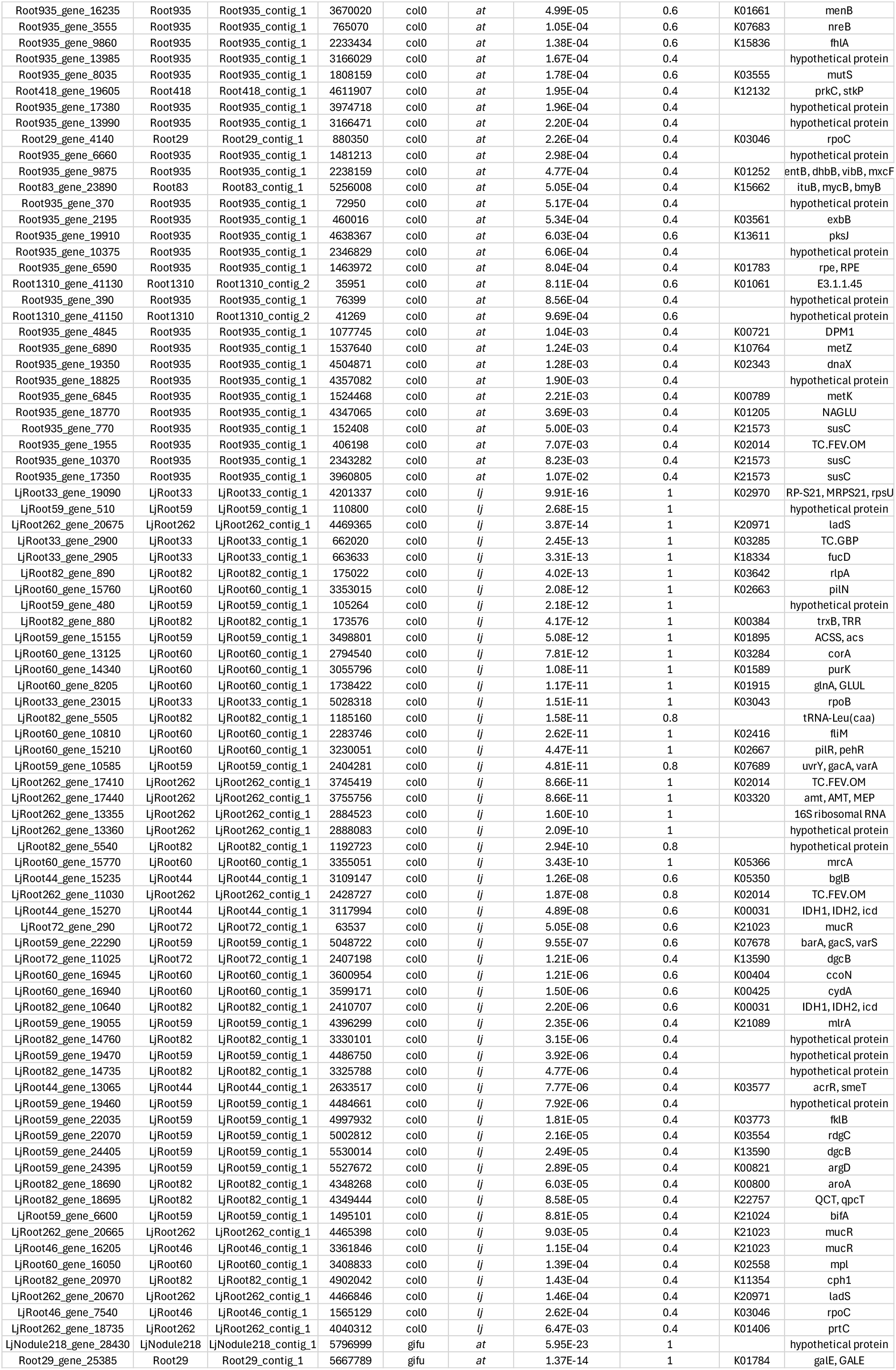
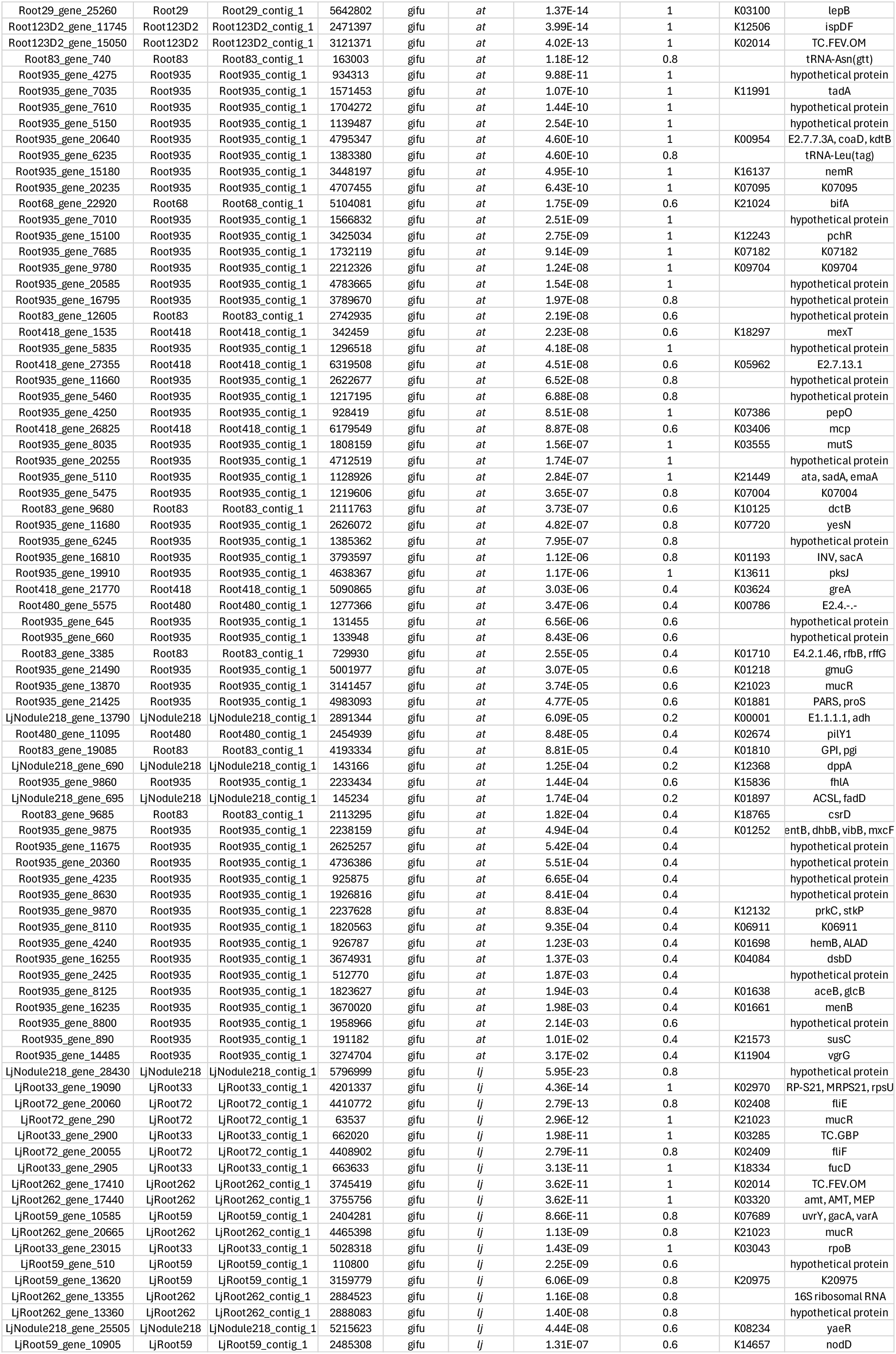
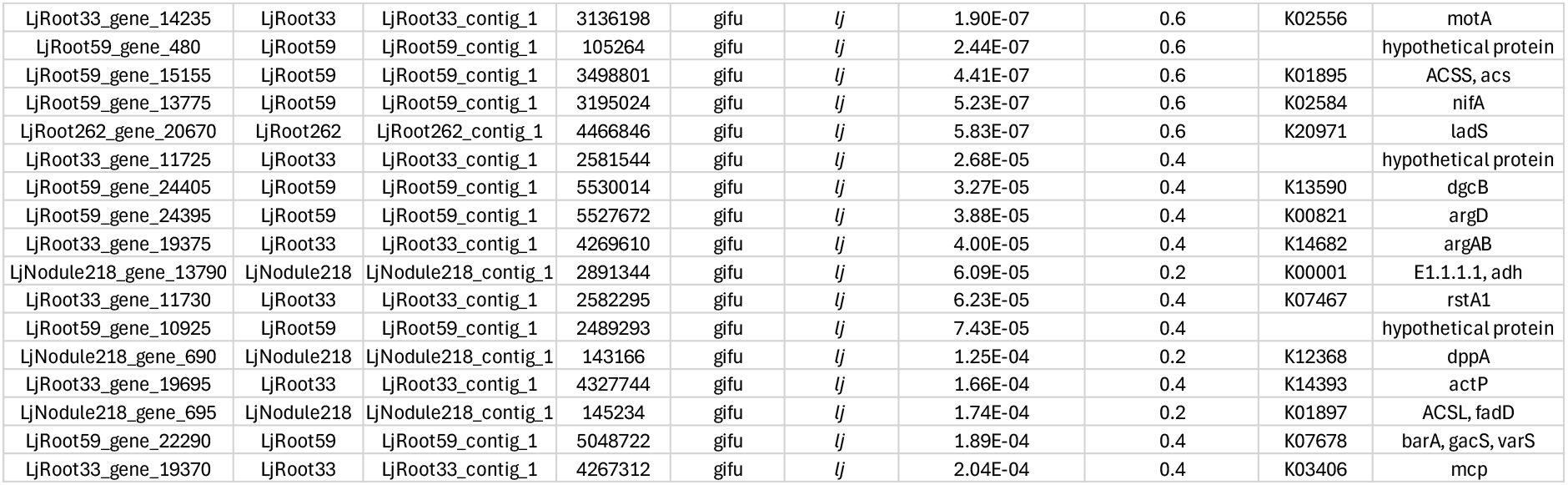
Genes exhibiting non-random patterns of mutation across replicate populations. The table lists the gene ID, strain, genomic coordinates, evolution host plant, and corresponding *P*-values of genes showing patterns of non-randomness (*n* = 259). Non-randomness was assessed at the ORF level using binomial tests, and *P*-values were adjusted for multiple testing using false discovery rate (FDR) control (*α* = 0.05). In addition, the table indicates the percentage of replicate populations in which mutations were detected for each ORF, as well as KEGG ontology identifiers.

**Table S3.**
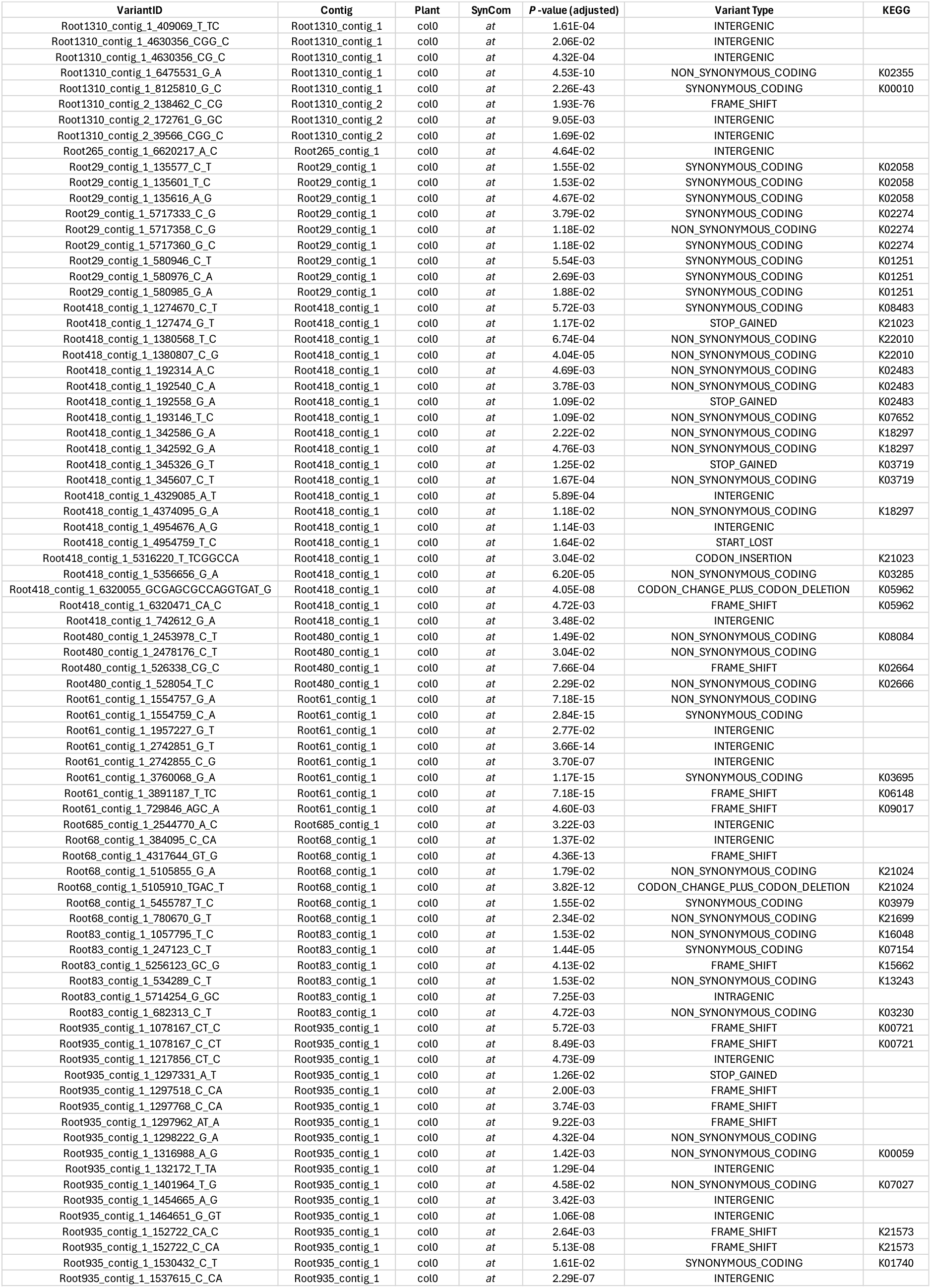
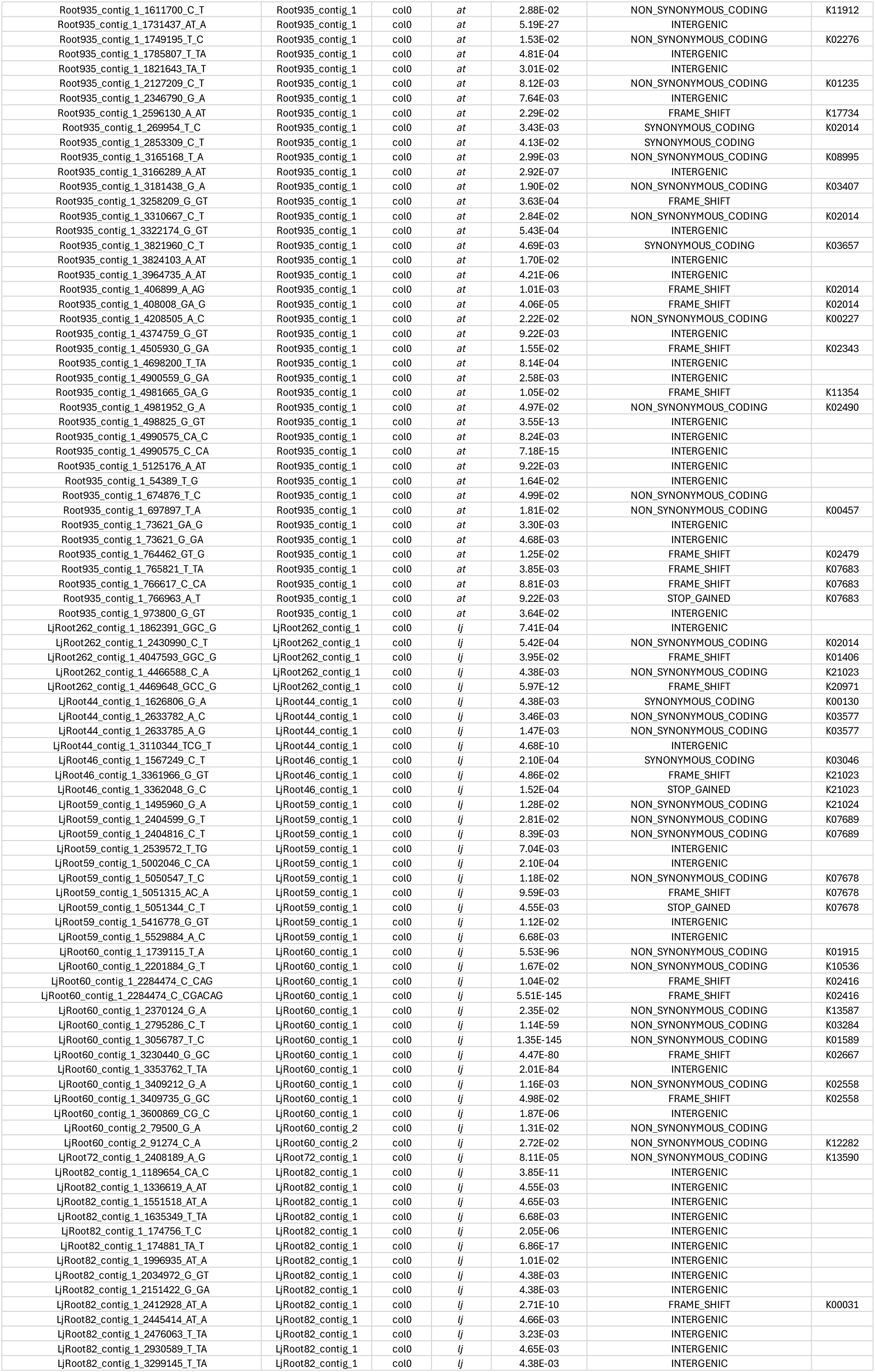
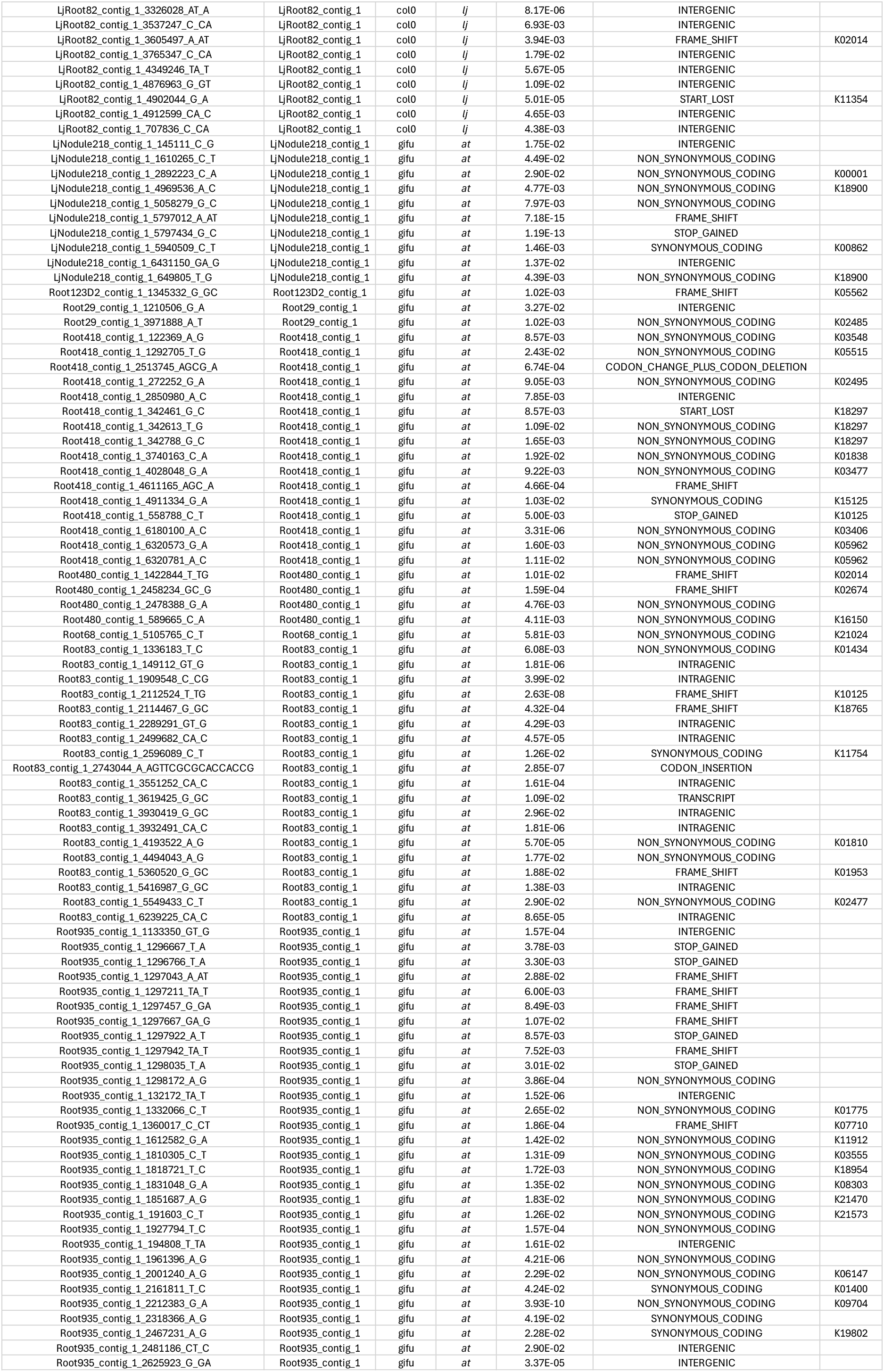
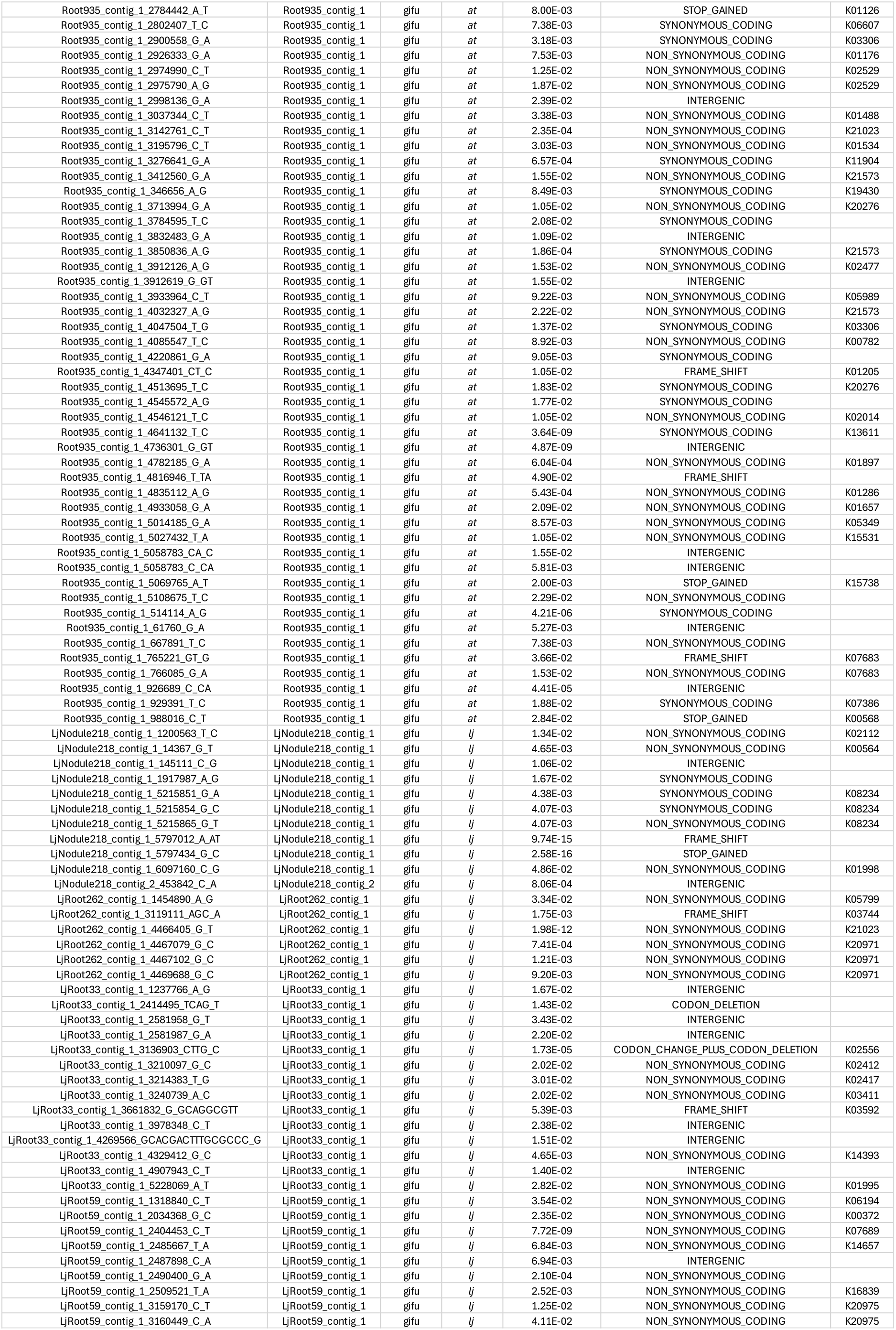
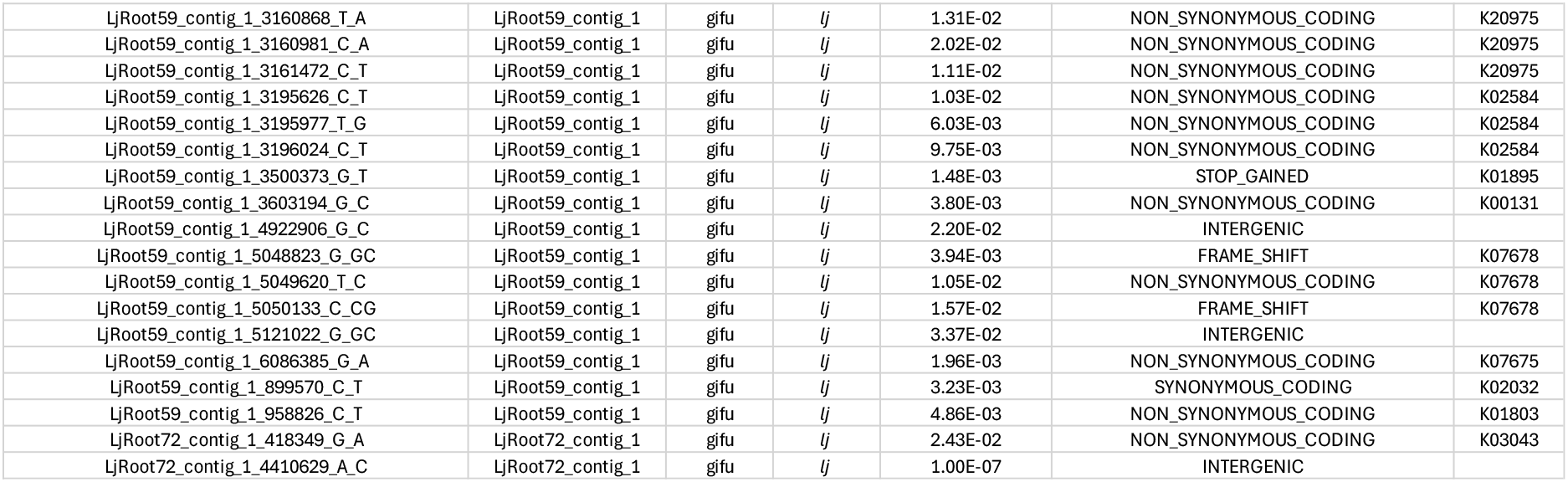
Genetic variants significantly enriched in SynCom populations evolved on *Arabidopsis* or *Lotus* roots. The table lists the variant ID, strain, genomic position, evolution host plant, and corresponding *P*-value (*n* = 372). Enrichment was assessed using generalized linear models (GLMs), and *P*-values were corrected for multiple testing by controlling the false discovery rate at (*α* = 0.05). KEGG ontologies are provided for mutations that affect coding regions.

**Table S4.** Structural genome rearrangements detected in the whole-genome sequencing of evolved isolates. The table lists the strain, genomic regions, and type of structural variant (SV; *n* = 12). In addition, the variation type (insertion or deletion), length, and corresponding evolution host are provided. For insertions, the likely origin of the inserted DNA fragment is indicated.

| Strain | Family | Contig | Location | Type | Length | Plant | SynCom | Condition | Insertion Origin |
| --- | --- | --- | --- | --- | --- | --- | --- | --- | --- |
| LjRoot262 | Sphingomonadaceae | LjRoot262_contig_1 | 2703013 | DEL | -108 | gifu | lj (gifu)-1 | native |  |
| LjRoot46 | Rhizobiaceae | LjRoot46_contig_5 | 50198 | INS | 1417 | gifu | lj (gifu)-1 | native | LjRoot46_contig_4 |
| LjRoot59 | Pseudomonadaceae | LjRoot59_contig_1 | 5734594 | DEL | -620 | gifu | lj (gifu)-1 | native |  |
| LjRoot59 | Pseudomonadaceae | LjRoot59_contig_1 | 5859699 | DEL | -596 | gifu | lj (gifu)-1 | native |  |
| LjRoot59 | Pseudomonadaceae | LjRoot59_contig_1 | 4453767 | DEL | -604 | gifu | lj (gifu)-1 | native |  |
| LjRoot60 | Xanthomonadaceae | LjRoot60_contig_1 | 3887682 | DEL | -247 | gifu | lj (gifu)-1 | native |  |
| LjRoot80 | Mycobacteriaceae | LjRoot80_contig_1 | 820540 | DEL | -702 | gifu | lj (gifu)-1 | native |  |
| LjRoot262 | Sphingomonadaceae | LjRoot262_contig_1 | 2703013 | DEL | -108 | col0 | lj (col0)-1 | non-native |  |
| LjRoot59 | Pseudomonadaceae | LjRoot59_contig_1 | 5859725 | DEL | -596 | col0 | lj (col0)-1 | non-native |  |
| LjRoot59 | Pseudomonadaceae | LjRoot59_contig_1 | 5734568 | DEL | -620 | col0 | lj (col0)-1 | non-native |  |
| LjRoot59 | Pseudomonadaceae | LjRoot59_contig_1 | 4453767 | DEL | -604 | col0 | lj (col0)-1 | non-native |  |
| LjRoot60 | Xanthomonadaceae | LjRoot60_contig_1 | 3887682 | DEL | -247 | col0 | lj (col0)-1 | non-native |  |

**Table S5.**
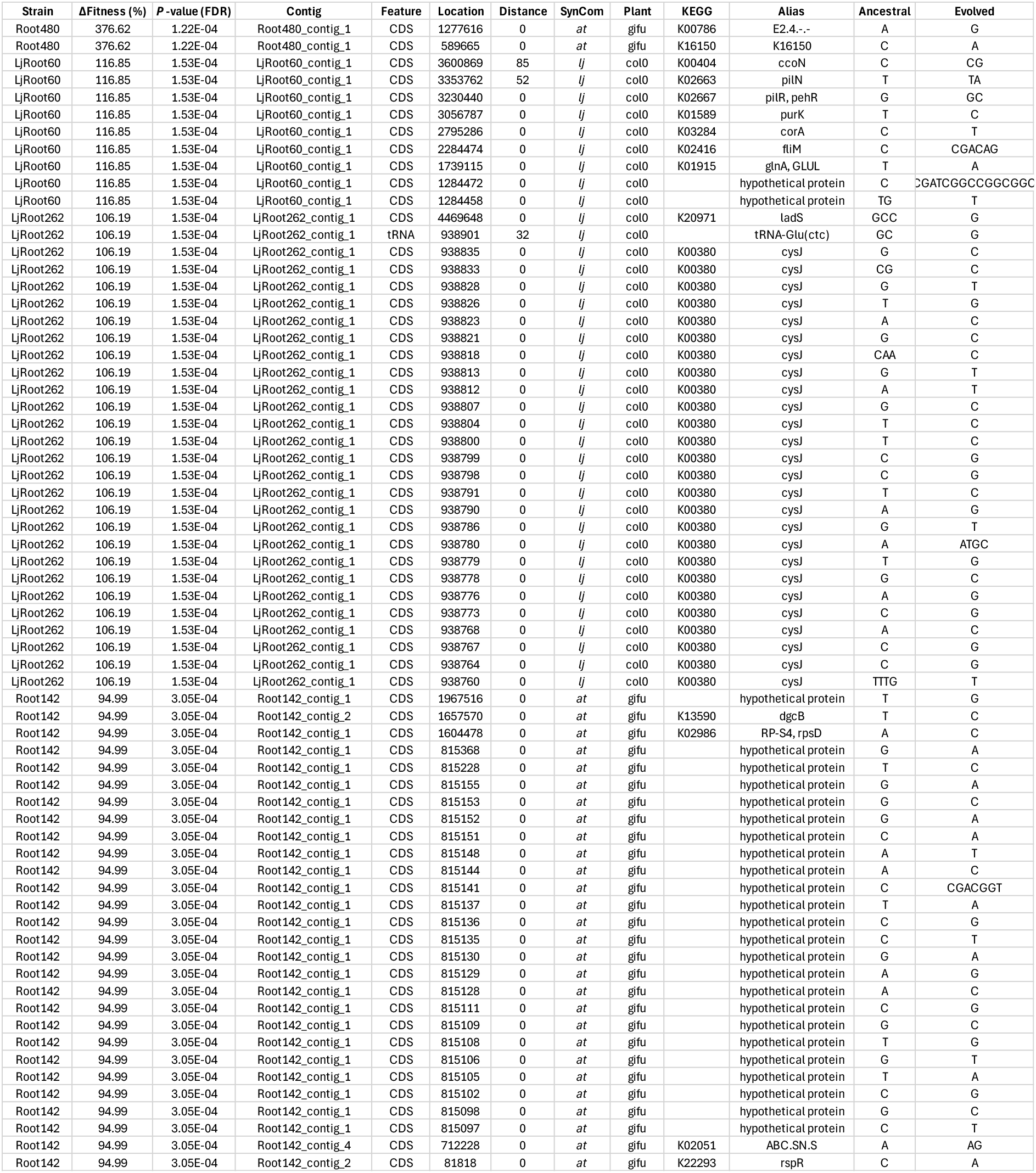
Mutations identified in highly adapted evolved isolates. For each variant (*n* = 67), the strain background, percent fitness change, *P*-value, genomic region, distance to the nearest downstream ORF, and corresponding evolution host are provided. For mutations within coding sequences, the distance was set to zero. The associated KEGG ontology and gene alias are listed.

**Table S6.** Dilution factors used for the culture collection of evolved isolates. For each reisolation, the experimental condition (SynCom and plant), plant generation, dilution factor and number of plates are provided.

| **Plant** | **SynCom** | **Generation** | **Dilution** | **#Plates** |
| --- | --- | --- | --- | --- |
| col0 | *at* | 1 | 18000 | 15 |
| col0 | *lj* | 1 | 18000 | 15 |
| gifu | *at* | 1 | 54000 | 15 |
| gifu | *lj* | 1 | 54000 | 15 |
| col0 | *at* | 3 | 18000 | 15 |
| col0 | *lj* | 3 | 18000 | 15 |
| gifu | *at* | 3 | 54000 | 15 |
| gifu | *lj* | 3 | 54000 | 15 |
| col0 | *at* | 5 | 18000 | 15 |
| col0 | *lj* | 5 | 18000 | 15 |
| gifu | *at* | 5 | 54000 | 15 |
| gifu | *lj* | 5 | 54000 | 15 |
| col0 | *at* | 7 | 36000 | 15 |
| col0 | *lj* | 7 | 36000 | 15 |
| gifu | *at* | 7 | 100000 | 15 |
| gifu | *lj* | 7 | 100000 | 15 |
| col0 | *at* | 9 | 56000 | 15 |
| col0 | *lj* | 9 | 56000 | 15 |
| gifu | *at* | 9 | 100000 | 15 |
| gifu | *lj* | 9 | 100000 | 15 |
| col0 | *at* | 11 | 56000 | 15 |
| col0 | *lj* | 11 | 56000 | 15 |
| gifu | *at* | 11 | 62500 | 15 |
| gifu | *lj* | 11 | 62500 | 15 |
| col0 | *at* | 13 | 54000 | 15 |
| col0 | *lj* | 13 | 54000 | 15 |
| gifu | *at* | 13 | 62000 | 15 |
| gifu | *lj* | 13 | 62000 | 15 |
| col0 | *at* | 15 | 54000 | 15 |
| col0 | *lj* | 15 | 54000 | 15 |
| gifu | *at* | 15 | 65000 | 15 |
| gifu | *lj* | 15 | 65000 | 15 |
| col0 | *at* | 16 | 54000 | 30 |
| col0 | *lj* | 16 | 54000 | 30 |
| gifu | *at* | 16 | 65000 | 30 |
| gifu | *lj* | 16 | 65000 | 30 |

**Table S7.** Summary of the dataset used for training the random forest classifier. The table reports the number of individual training instances for each strain and for the TY-medium control, comprising a total of 35 classes used to train the classifier on fluorescence spectral data.

| **Strain** | **Replicates** |
| --- | --- |
| Root131 | 144 |
| LjRoot44 | 144 |
| Root935 | 144 |
| Root61 | 144 |
| LjRoot60 | 144 |
| LjRoot303 | 144 |
| Root77 | 144 |
| Root83 | 144 |
| LjRoot33 | 144 |
| LjRoot80 | 144 |
| Root123D2 | 144 |
| LjRoot17 | 144 |
| LjRoot24 | 216 |
| Root1310 | 216 |
| LjRoot5 | 216 |
| Root142 | 216 |
| Root685 | 216 |
| LjRoot262 | 216 |
| LjRoot52 | 216 |
| LjRoot59 | 216 |
| LjRoot82 | 216 |
| LjRoot222 | 216 |
| Root720 | 216 |
| Root265 | 216 |
| LjRoot46 | 216 |
| LjNodule218 | 216 |
| LjRoot72 | 216 |
| Root68 | 216 |
| Root480 | 216 |
| Root418 | 216 |
| LjRoot1 | 216 |
| Root29 | 216 |
| Root101 | 216 |
| Root695 | 216 |
| medium | 432 |

**Table S8.** Evolved isolates used for reconstitution experiments. Reisolated strains from the evolution experiment are listed along with their respective metadata (*n* = 34). Columns include a unique identifier, original strain name, taxonomic family, SynCom, evolution host (i.e., the host on which the strain evolved during the experiment), the experimental evolution condition, and the competition experiments in which the strain was used.

| ID (Reisolation) | Strain | Family | SynCom | Plant | Condition | Experiment |
| --- | --- | --- | --- | --- | --- | --- |
| 53 | LjNodule218 | Phyllobacteriaceae | *lj* | gifu | native | exp_1 |
| 2-23 | LjNodule218 | Phyllobacteriaceae | *at* | gifu | non-native | exp_1 |
| 62 | LjRoot17 | Caulobacteraceae | *lj* | gifu | native | exp_1 |
| 80 | LjRoot222 | Hyphomicrobiaceae | *lj* | gifu | native | exp_1 |
| 2-43 | LjRoot222 | Hyphomicrobiaceae | *lj* | gifu | native | exp_1 |
| 2-44 | LjRoot262 | Sphingomonadaceae | *lj* | gifu | native | exp_1 |
| 2-3 | LjRoot262 | Sphingomonadaceae | *lj* | col0 | non-native | exp_1 |
| 72 | LjRoot33 | Oxalobacteraceae | *lj* | gifu | native | exp_1 |
| 2-43 | LjRoot33 | Oxalobacteraceae | *lj* | gifu | native | exp_1 |
| 74 | LjRoot44 | Microbacteriaceae | *lj* | gifu | native | exp_1 |
| 167 | LjRoot44 | Microbacteriaceae | *lj* | col0 | non-native | exp_1 |
| 2-45 | LjRoot46 | Rhizobiaceae | *lj* | gifu | native | exp_1 |
| 81 | LjRoot59 | Pseudomonadaceae | *lj* | gifu | native | exp_1 |
| 140 | LjRoot59 | Pseudomonadaceae | *lj* | col0 | non-native | exp_1 |
| 72 | LjRoot60 | Xanthomonadaceae | *lj* | gifu | native | exp_1 |
| 2-94 | LjRoot60 | Xanthomonadaceae | *lj* | col0 | non-native | exp_1 |
| 5 | LjRoot60 | Xanthomonadaceae | *lj* | col0 | non-native | exp_1 |
| 54 | LjRoot72 | Comamonadaceae | *lj* | gifu | native | exp_1 |
| 138 | LjRoot72 | Comamonadaceae | *lj* | col0 | non-native | exp_1 |
| 61 | LjRoot80 | Mycobacteriaceae | *lj* | gifu | native | exp_1 |
| 2-68 | LjRoot80 | Mycobacteriaceae | *at* | col0 | native | exp_1 |
| 71 | LjRoot82 | Flavobacteriaceae | *lj* | gifu | native | exp_1 |
| 2-94 | LjRoot82 | Flavobacteriaceae | *lj* | col0 | non-native | exp_1 |
| 90 | Root142 | Rhizobiaceae | *at* | col0 | native | exp_1 |
| 44 | Root142 | Rhizobiaceae | *at* | gifu | non-native | exp_1 |
| 2-74 | Root265 | Mycobacteriaceae | *at* | col0 | native | exp_1 |
| 21 | Root480 | Xanthomonadaceae | *at* | gifu | non-native | exp_1 |
| 93 | Root61 | Microbacteriaceae | *at* | col0 | native | exp_1 |
| 2-34 | Root61 | Microbacteriaceae | *at* | gifu | non-native | exp_1 |
| 2-68 | Root685 | Hyphomicrobiaceae | *at* | col0 | native | exp_1 |
| 11 | Root77 | Caulobacteraceae | *at* | gifu | non-native | exp_1 |
| 2-77 | Root935 | Flavobacteriaceae | *at* | col0 | native | exp_1 |
| 134 | Root935 | Flavobacteriaceae | *lj* | col0 | non-native | exp_1 |
| 18 | Root935 | Flavobacteriaceae | *at* | gifu | non-native | exp_1 |

